# Neuronal Autophagy Failure Drives α-Synuclein Transfer to Microglia to Outsource Aggregate Clearance

**DOI:** 10.1101/2024.04.19.590207

**Authors:** Ranabir Chakraborty, Philippa Samella, Veronica Testa, Jara Montero-Muñoz, Takashi Nonaka, Masato Hasegawa, Antonella Consiglio, Chiara Zurzolo

## Abstract

Tunneling nanotubes (TNTs) play a crucial role in intercellular communication, enabling a dynamic network for the transfer of molecular cargo over long distances between connected cells. Previous studies have demonstrated efficient, directional transfer of *α*-Synuclein (*α*-Syn) aggregates from neurons to microglia, with endosomal trafficking and lysosomal processing identified as the primary events following α-Syn internalization. Using human neuronal and microglial cell lines, we found that microglia exhibit higher lysosomal turnover, particularly through lysophagy, whereas neuronal lysosomes display compromised degradative capacity and impaired autophagic flux. This deficiency results in less efficient degradation of aggregates in neurons. Moreover, perturbation of autophagy enhances TNT-mediated transfer of aggregate from neuronal cells to microglia. In contrast, microglia co-cultured with *α*-Syn-containing neurons upregulate autophagy flux, enabling efficient degradation of the transferred aggregates. These findings were further validated using human induced pluripotent stem cells (hiPSC)-derived neurons and microglia. Overall, our study highlights the distinct responses of neurons and microglia to *α*-Syn aggregates and identifies dysfunctional autophagy in neurons as a key driver of the preferential and directional transfer of aggregates to microglia.

## Introduction

Parkinson’s Disease (PD) is the second most common neurodegenerative disease, the worldwide burden of which has doubled between 1990 and 2016 (Dorsey *et al*, 2018). One of the major genetic associations of sporadic PD, and the first to be discovered, is *SNCA*, which encodes a cytosolic protein *α*-Synuclein (*α*-Syn). Both mutations in *SNCA,* as well as elevated levels of the protein (due to duplication and triplication of the gene), can give rise to a pathological state (Polymeropoulos *et al*, 1997; Singleton *et al*, 2003; Miller *et al*, 2004).

A common pathological hallmark of several neurodegenerative diseases, including PD, is disrupted proteostasis. Under physiological conditions, quality control pathways in cells allow for efficient clearing of misfolded proteins, thereby preventing their accumulation and/or aggregation (Balch *et al*, 2008). Autophagy, a key proteostasis pathway, orchestrates the degradation of obsolete intracellular components, including organelles and aggregated proteins, via lysosomal machinery. Several reports highlight compromised macroautophagy (henceforth referred to as “autophagy”) or chaperone-mediated autophagy (CMA) in PD: impaired autophagosome formation (Winslow *et al*, 2010), dysregulated mitophagy due to mutations in genes encoding PTEN-induced putative kinase 1 (*PINK1*) and Parkin RBR E3 ubiquitin-protein ligase (*PARKIN*) (Valente *et al*, 2004; Kitada *et al*, 1998), decreased CMA caused by mutations in leucine-rich repeat kinase 2 (*LRRK2*) or *SNCA* (Orenstein *et al*, 2013; Alvarez-Erviti *et al*, 2010), and impaired lysosomal activity due to mutations in glucocerebrosidase-encoding gene *GBA1* (Sardi *et al*, 2011). Besides genetic mutations, *in vitro* overexpression of *α*-Syn in human mesencephalic neuronal (LUHMES) cells has also been reported to impair autophagy by preventing the fusion of autophagosomes with lysosomes (Tang *et al*, 2021). Additionally, overexpression of *α*-Syn in human neuroblastoma (SH-SY5Y) cells impairs starvation-induced autophagy (Nascimento *et al*, 2020). Such dysregulation prompts cells to seek alternative mechanisms to restore homeostasis, failure of which may culminate in apoptosis. Of note, the accumulation of intracellular α-Syn aggregates resulting from dysregulated autophagy has been associated with their secretion via exosomes and membrane shedding (Alvarez-Erviti *et al*, 2011; Danzer *et al*, 2012; Lee *et al*, 2013; Poehler *et al*, 2014). Moreover, recent findings elucidate the role of microglia in alleviating neuronal α-Syn burden through a process termed "synucleinphagy," wherein microglia engulf aggregates released by neurons and target them towards autophagy-mediated clearance (Choi *et al*, 2020). However, whether α-Syn aggregates have a different impact in lysosomal and autophagic pathways in neuronal versus microglial cells, and whether microglia modulate proteostasis in response to neuronal *α*-Syn burden, remain unknown. Since their discovery in 2004 (Rustom *et al*, 2004), tunneling nanotubes (TNTs) have emerged as pivotal conduits for contact-dependent cell-to-cell communication that mediate the exchange of various molecular cargo, from small ions to larger organelles (Cordero Cervantes & Zurzolo, 2021). Moreover, these membranous, F-actin-rich structures facilitate the movement of neurodegenerative disease-associated protein aggregates such as α-Syn, tau and Amyloid-β (Chakraborty *et al*, 2023a). α-Syn aggregates exploit TNTs to traverse between diverse cell types, including neuronal precursor cells (NPCs) (Grudina *et al*, 2019), neurons (Senol *et al*, 2021; Abounit *et al*, 2016), astrocytes (Rostami *et al*, 2017; Loria *et al*, 2017), and microglia (Scheiblich *et al*, 2021; Chakraborty *et al*, 2023b). Even though previous studies have demonstrated the augmentation of homotypic connections between neuronal cells, and microglial cells by α-Syn aggregates (Senol *et al*, 2021; Chakraborty *et al*, 2023b), the mechanism underlying their preferential directional transfer from neuronal cells to microglia remains enigmatic.

In this study, we investigate the distinct subcellular dynamics of lysosomal processing abilities in neuronal and microglial cells, both in the presence and absence of α-Syn aggregates. We also examine the non-cell autonomous influence of aggregate-burdened neuronal cells on autophagy regulation in microglia. Our findings uncover a fundamental imbalance in lysosomal function and autophagic activity between neurons and microglia which drives the preferential transfer of α-Syn aggregates from neurons towards microglia. They advance our understanding of how organellar dysfunction shapes neuron–microglia crosstalk in the context of pathological protein aggregation.

## Results

### *α*-Syn localization to lysosomes in neuronal cells and microglia

To assess the sub-cellular localization of *α*-Syn pre-formed fibrils, we challenged both SH-SY5Y neuronal and HMC3 microglial cells with aggregates for different time durations, ranging from 1 hour to 16 hours, and performed immunostaining against lysosome-associated membrane protein 1 (LAMP1) (**Fig. 1A**). We observed a time-dependent increase in the presence of *α*-Syn aggregates with lysosomes, a phenotype similar for both the cell types. However, the extent of lysosomal association with aggregates differed markedly between neurons and microglia (**Fig. 1B, C**). In neuronal cells, following *α*-Syn exposure for 16 hours, an average of 67.58% of lysosomes overlapped with *α*-Syn, whereas only 29.30% of microglial lysosomes localized with aggregates at that time point. The increase in lysosomal localization over time was steeper in neuronal cells than in microglia, as reflected by the slope of the linear regression fit (0.04249 vs. 0.01620, respectively). Structured illumination super-resolution microscopy further confirmed greater co-localization of α-Syn aggregates with neuronal lysosomes than with microglial lysosomes after 16 hours of exposure (**Fig. 1D, E**; *cyan asterisks*). Taken together, these results highlight cell type–specific differences in aggregate–lysosome association, suggesting a potential disparity in the impact of α-Syn aggregates on lysosomal function.

**Figure 1.**
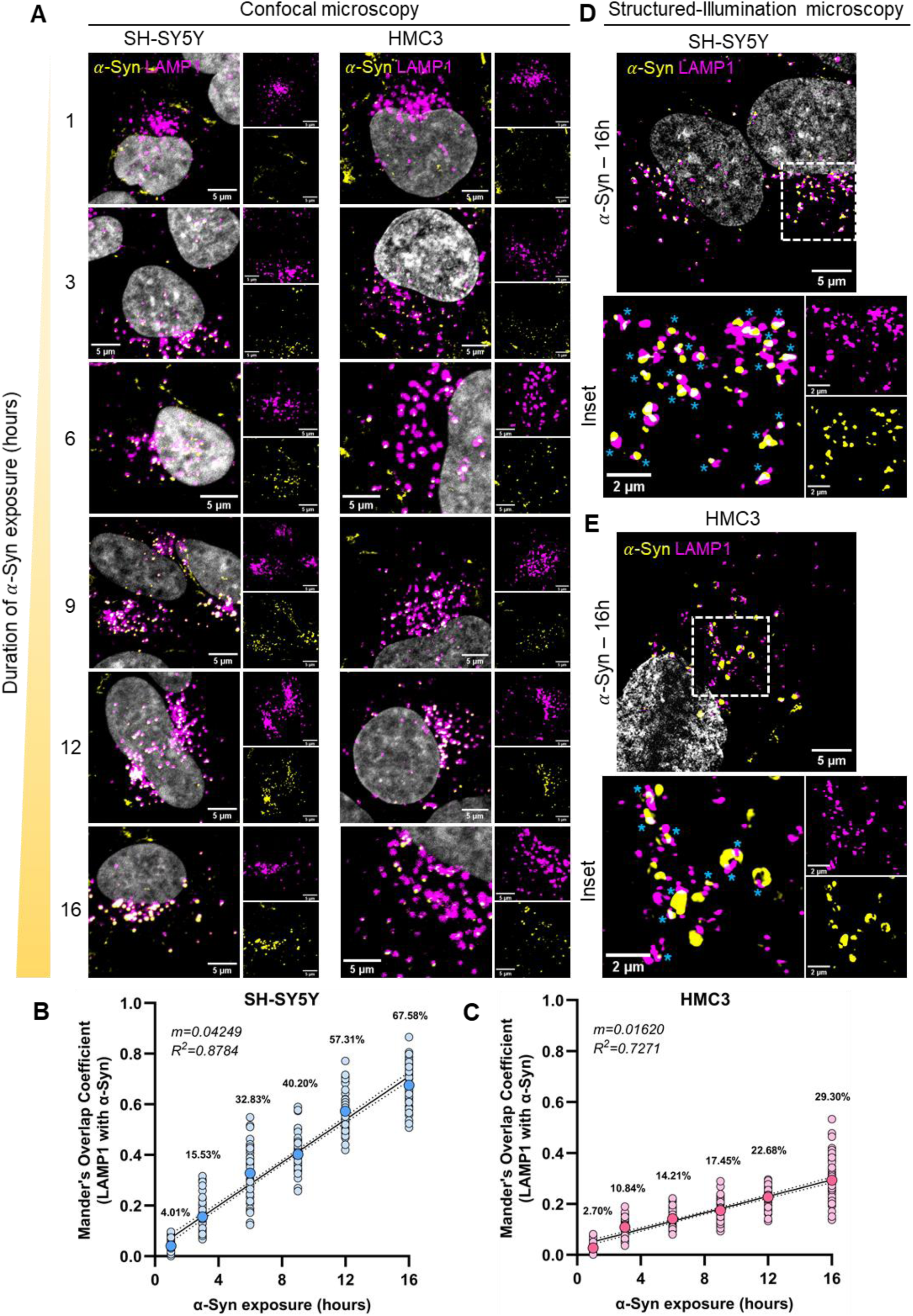
Temporal analysis *α*-Syn localization in neuronal cells and microglia. (**A**) Representative confocal images of *α*-Syn (pseudo-colored yellow) localization with lysosomes (pseudo-colored magenta) at different (indicated) time points of aggregate exposure of SH-SY5Y neuronal cells (left panels) and HMC3 microglia (right panels). (**B and C**) Quantification of co-localization between lysosomes and aggregates (Mander’s overlap coefficient – fraction of LAMP1 with *α*-Syn) at different time points of aggregate exposure of neuronal cells (**B**) and microglia (**C**). N=3 independent experiment, n=50 cells per group. Larger, darker filled circle depicts the mean, also mentioned as percentages within the graphs. Black line denotes the simple linear regression fit, dotted lines representing the 95% confidence intervals. (**D and E**) Structured-Illumination microscopy images of neuronal cells (**D**) and microglia (**E**) exposed to aggregates for 16 hours show localization of *α*-Syn to lysosomes. Images are pseudo-colored – magenta for LAMP1-647, yellow for *α*-Syn488, and gray for DAPI. Cyan asterisks denote areas of *α*-Syn – lysosome colocalization.

### *α*-Syn aggregates differentially affect lysosomes in neuronal cells and microglia

To assess the impact of α-Syn aggregates on lysosomal function, we evaluated both lysosomal degradative capacity (**Fig. 2**) and motility (**Fig. S2**). Cells were first incubated with dextran-647 (10 kDa) for 3 hours (pulse phase), followed by exposure to α-Syn aggregates for 16 hours (chase phase), during which dextran traffics from early endosomes and predominantly localizes to lysosomes (Barral *et al*, 2022). Two hours before the end of the chase period, cells were incubated with dye-quenched bovine serum albumin (DQ-BSA), which is internalized and trafficked to lysosomes (**Fig. 2A**).Catalytic cleavage of DQ-BSA-dye results in substantial increase in fluorescence (Marwaha & Sharma, 2017), thereby allowing to probe the functionality of lysosomes. Exposure to *α*-Syn aggregates markedly reduces the proportion of degradative lysosomes (DQ-BSA^+^Dextran^+^; indicated by cyan asterisks in representative images) per cell in both the cell types (**Fig. 2B-D**). However, the reduction was more pronounced in neuronal cells — dropping from 81.86% under control conditions to 45.01% following aggregate treatment— compared to a smaller decline in microglia (from 83.59% to 75.63%). These data were further supported by quantification of DQ-BSA fluorescence intensity in dextran+ lysosomes, a read-out of lysosomal degradative ability, which revealed a significant reduction in neuronal cells but not in microglia (**Fig. 2E-F**).

**Figure 2.**
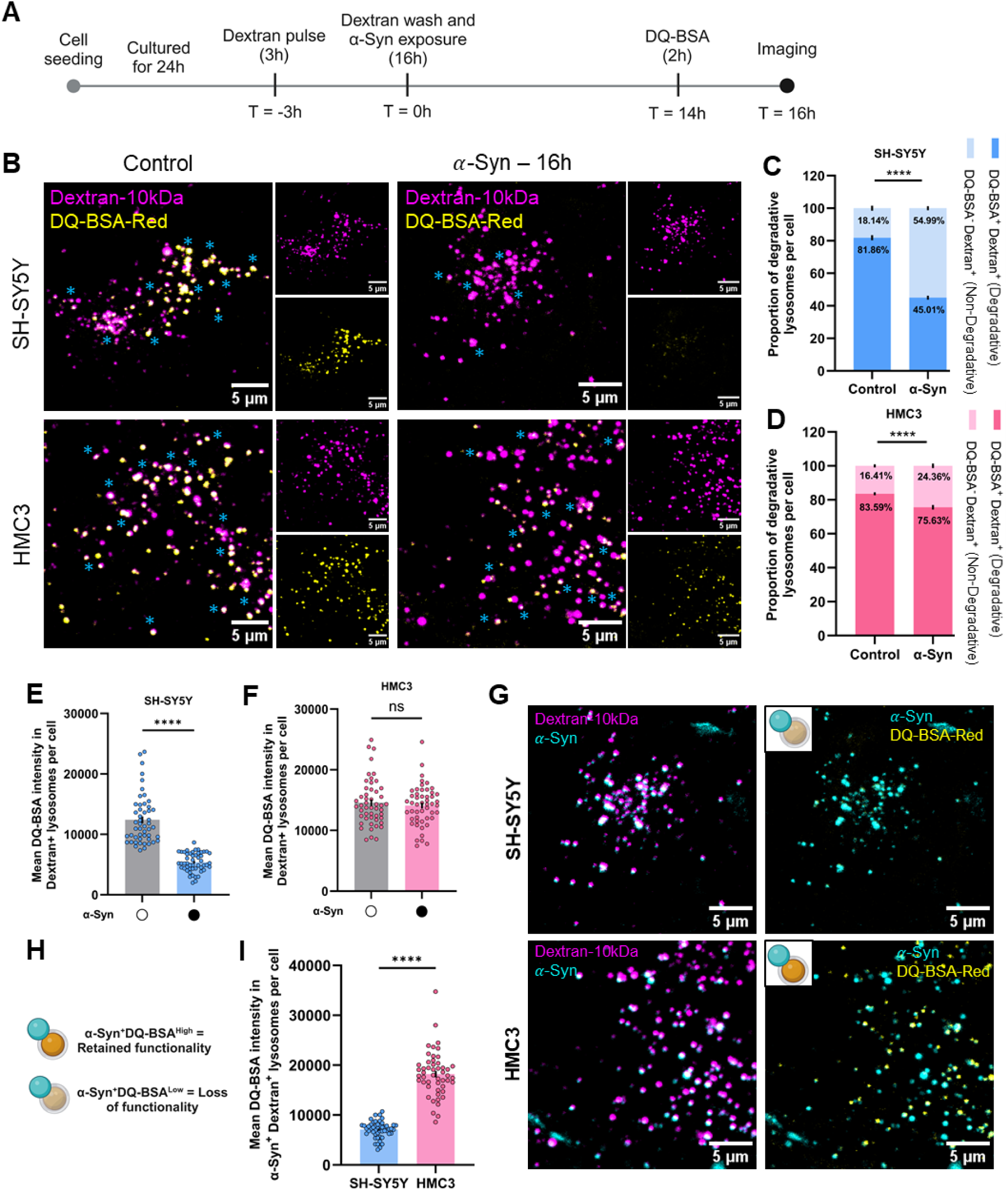
Effect of *α*-Syn aggregates on lysosomal degradative ability. (**A**) Schematic representation of the experimental design to assess for degradative lysosomes. (**B**) Representative images of neuronal and microglial cells pulse-chased for Dextran647-10kDa (magenta) and DQ-BSA red (yellow) in SH-SY5Y (upper panels) and HMC3 (lower panels) cells. (**C and D**) Quantification of the proportion of degradative lysosomes per cell as a percentage of total number of lysosomes for neuronal cells (**C**) and microglia (**D**). N=3 independent experiments, n=50 cells per group. Mean values are mentioned within the graph, error bars representing SEM. Statistical significance was analyzed using Two-Way ANOVA with uncorrected Fisher’s LSD multiple comparison. ****p<0.0001. (**E and F**) Quantification of mean DQ-BSA fluorescence intensities in Dextran+ neuronal (**E**) and microglial (**F**) lysosomes (lysosomal degradative ability per cell). N=3 independent experiments, n=50 cells per group. Statistical significance was analyzed using two-tailed Mann-Whitney U test. ns: non-significant (p=0.8611), ****p<0.0001. (**G**) Representative dual-channel images of neuronal cells (left panels) and microglia (right panels) treated with *α*-Syn488 (cyan). (**H and I**) Schematic representation (**H**) of the quantification (**I**) of degradative ability of lysosomes associated with *α*-Syn. N=3 independent experiments, n=50 cells per group. Statistical significance was analyzed using two-tailed Mann-Whitney U test. ****p<0.0001.

We also observed a differential effect on the functionality of lysosomes localized with aggregates (α-Syn⁺/Dextran⁺). In neuronal cells, these lysosomes exhibited reduced DQ-BSA fluorescence, indicating compromised degradative function, whereas in microglia, lysosomes with α-Syn aggregates largely retained their functionality (**Fig. 2G–I**). These findings highlight a cell type–specific vulnerability, revealing that neuronal lysosomes are more functionally sensitive to α-Syn aggregates at the single-organelle level.

Lysosomal functionality depends on their membrane integrity. Lysosome membrane permeabilization (LMP) has been reported in both PD, and cells exposed to extracellular *α*-Syn aggregates (Senol *et al*, 2021; Vila *et al*, 2011; Dehay *et al*, 2013). To prevent such ruptures, different ESCRT proteins are known to be recruited on damaged lysosomes to facilitate membrane repair (Radulovic *et al*, 2018). To examine occurrence of LMP and lysosomal membrane repair in neuronal and microglial cells upon *α*-Syn treatment, we performed immunostaining against Galectin-3, a marker of endo-lysosomal rupture (**Fig. S1A**) and the ESCRT-III associated factor IST-1 (**Fig. S1B**). We observed increased LMP upon aggregate exposure in both neuronalcells compared to microglial cells (**Fig. S1C, D**). However, the extent of LMP was more than 3 folds higher in neuronal cells compared to microglia [2.83% (of total lysosomes) in neuronal cells versus 0.78% in microglia]. Accordingly, the need for membrane repair was also different between the two cell types (2.82% of neuronal lysosomes being IST-1+, versus 1.13% for microglia) (**Fig. S1E, F**). Taken together, these results demonstrate higher sensitivity of neuronal lysosomes to *α*-Syn aggregates compared to those in microglia, highlighting the relative resilience of the latter.

Given the importance of lysosomal trafficking in regulating their positioning, and consequently their function (Scerra *et al*, 2022), we assessed whether aggregate association influenced lysosomal motility. Using lysotracker green, we tracked individual lysosomes over 2 minutes in control cells and in cells treated with aggregates for 16h, comparing lysosomes containing aggregates (+*α*-Syn) or not (-*α*-Syn) (**Fig. S2A-D; and supplementary videos 1-6**). The presence of aggregates on lysosomes significantly reduced both their mean travel distance (**Fig. S2E and G**) and their mean velocities (**Fig. S2F and H**) in both neuronal and microglia cells. Additionally, aggregate-bearing lysosomes were more likely to be classified as less motile—defined arbitrarily as traveling ≤15 μm, based on the median track length of control neuronal lysosomes (16.837 μm)— in both cell types (**Fig. S2I, J**). Notably, the proportion of less motile lysosomes was higher in neuronal cells than in microglia under both control and α-Syn–treated conditions (34% vs. 14% in controls, and 54% vs. 30% in aggregate-containing lysosomes). A cumulative frequency distribution further reported a higher number of lysosomes with reduced distance travelled upon association with *α*-Syn aggregates (**Fig. S2K, L;** arrow depicting a leftward shift in the peak).

Taken together, these data indicate that neuronal lysosomes are more susceptible to the presence of aggregates than those in microglia, resulting in compromised degradative capability. Interestingly, the movement of individual lysosomes appears to be similarly affected by the presence of aggregates in both neuronal and microglial cells. These observations are consistent with the greater localization of aggregates to lysosomes in neuronal cells (**Fig. 1**), ultimately contributing to the observed reduction in lysosomal functionality.

### Lysophagy and lysosome biogenesis are elevated in microglia

Building on our observation of higher aggregate localization with, and consequent damage to, lysosomes in neuronal cells, we hypothesized that the resilience of microglial lysosomes could be attributed to a more efficient clearance and biogenesis of these organelles in microglia compared to neuronal cells. Since p62 has been reported to be recruited to damaged lysosomes to facilitate their clearance via lysophagy (Gallagher & Holzbaur, 2023), we assessed the extent of colocalization of LAMP1-positive lysosomes with p62 under different conditions. To distinguish lysosomes destined for lysophagy from autolysosomes (which also show p62-LAMP1 co-localization), we conducted this experiment in the presence of bafilomycin A1, which prevents autophagosome-lysosome fusion, thereby allowing the identification of lysosomes marked for lysophagy by p62 (**Fig. 3A**, cyan dotted circles in inset images). As expected, the extent of lysosomes undergoing lysophagy was significantly increased upon LLOMe exposure in both neuronal and microglial cells (**Fig. 3B-C**). Similarly, exposure to aggregates for 16h also increased lysophagy in both cell types. In neuronal cells, lysophagy increased from a basal level of 5.95% to 21.16% upon aggregate exposure (a 15.21% increase) (**Fig. 3B).** In microglia, lysophagy rose from a basal level of 25.63% to 49.20% (a 23.57% increase) **(Fig. 3C)**. Notably, lysophagy was significantly higher in microglia compared to neuronal cells, both at a basal level, and following aggregate exposure (**Fig. 3D**). These results point towards an enhanced ability of microglia to clear damaged lysosomes and restore proteostasis, a process that appears to be less efficient in neuronal cells.

**Figure 3.**
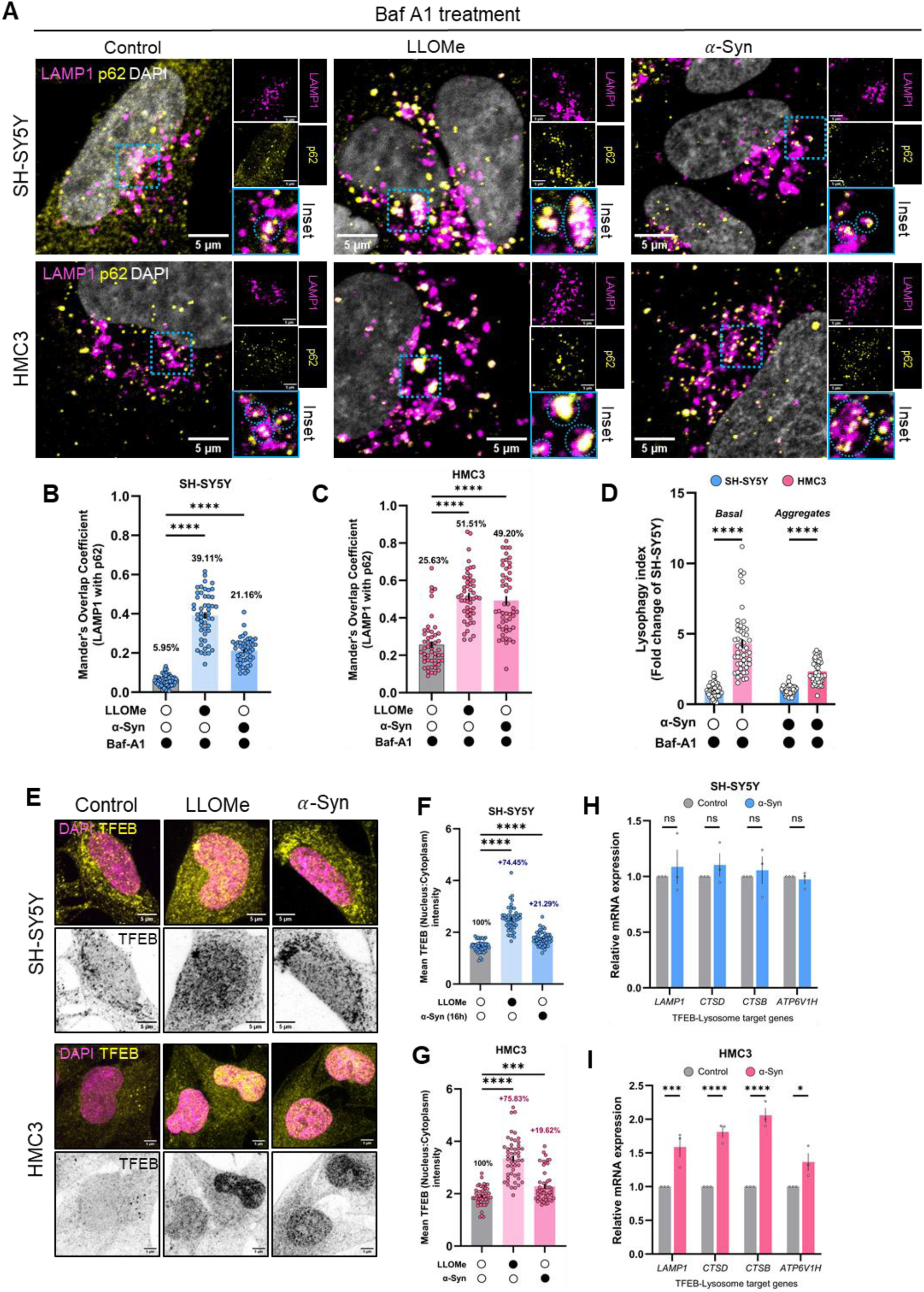
Effect of aggregates on lysophagy and lysosome biogenesis. **(A)** Representative images of SH-SY5Y cells (upper panels) and microglia (bottom panels) immunostained for LAMP1 (magenta) and p62 (yellow) in the presence of bafilomycin A1. (**B**) Quantification of the extent of co-localization of LAMP1-positive structures with p62 as a readout for lysophagy in SH-SY5Y cells. Co-localization is represented as Mander’s Overlap Coefficient of the extent of LAMP1 overlap with p62. Mean percentage of co-localization is mentioned within the graph. N=3 independent experiments, n=50 cells per group. Statistical significance was analyzed using Brown-Forsythe and Welch One-Way ANOVA with Dunnett’s T3 multiple comparison. ****p<0.0001. (**C**) Same quantification as in (**B**), but in HMC3 cells. Error bars represent SEM. Statistical significance was analyzed using Kruskal-Wallis test with Dunn’s multiple comparison. ****p<0.0001. (**D**) Comparative analysis of the fold-difference in lysophagy induction between neuronal and microglial cells under basal conditions or upon aggregate-exposure. Statistical significance was analyzed using Two-Way ANOVA with Šídák’s multiple comparison. ****p<0.0001. (**E**) Representative images of TFEB immunostaining (yellow, and gray-invert) in SH-SY5Y (upper panels) and HMC3 (bottom panels) cells in different conditions. (**F**) Quantification of the extent of TFEB translocation to the nucleus (mean TFEB intensity: nucleus-to-cytoplasmic ratio) in SH-SY5Y cells. Percentage increase relative to control (100%) is mentioned within the graph. (**G**) Same quantification as in (**F**), but in HMC3 cells. (**H**) Relative expression level of TFEB-lysosomal target genes in SH-SY5Y cells. N=3 biological replicates. Error bars represent SEM. Statistical significance was analyzed using Two-Way ANOVA with Šídák’s multiple comparison (comparison between control and *α*-Syn-exposed cells). ns: non-significant (p=0.9157 for *LAMP1*, 0.8492 for *CTSD*, 0.9822 for *CTSB*, and 0.9989 for *ATP6V1H*). (**I**) Relative expression level of TFEB-lysosomal target genes in HMC3 cells. N=3 biological replicates. Error bars represent SEM. Statistical significance was analyzed using Two-Way ANOVA with Šídák’s multiple comparison (comparison between control and *α*-Syn-exposed cells). *p=0.0325, ***p=0.0007, ****p<0.0001.

As part of lysophagic flux, the clearance of organelles necessitates homeostatic compensation through lysosomal biogenesis. Transcription factor EB (TFEB) is a major regulator of autophagy and lysosome-associated genes (Settembre *et al*, 2011). We performed immunostaining against TFEB in both neuronal cells and microglia, and assessed the extent of its nuclear translocation. Consistent with our observation of increased lysophagy within 16h of aggregate exposure, nuclear translocation of TFEB also increased in both the cell types (**Fig. 3E-G**). To further evaluate TFEB activity, we performed RT-PCR to quantify the mRNA levels of TFEB-regulated lysosomal target genes. Upon aggregate exposure, microglial gene expression of the lysosomal protein (*LAMP1*), hydrolases (*CTSD* and *CTSB*), and a subunit of the vacuolar ATPase (*ATP6V1H*) significantly increased relative to control, a phenotype not observed in neuronal cells (**Fig. 3H, I**). Taken together, these results suggest an increase in lysophagy upon exposure to *α*-Syn aggregates in both the cell types, with microglia being relatively more efficient in lysosomal turnover and biogenesis compared to neuronal cells.

### *α*-Syn aggregates differentially affect autophagic flux in neuronal cells and microglia

To understand the consequence of the differential lysosomal response to *α*-Syn between neuronal cells and microglia, we investigated whether aggregates had any impact on autophagy, given its reliance on functional lysosomes. A robust method for measuring autophagy flux is to assess the conversion of LC3B from its non-lipidated (LC3B-I) form to its lipidated form (LC3B-II), which is associated with autophagosomes (**Fig. 4A**). As expected, bafilomycin A1 treatment of cells led to an accumulation of LC3B-II in both neuronal and microglial cells (**Fig. 4A-C**). Interestingly, *α*-Syn exposure of neuronal cells led to a more-than 2-fold decline in LC3B-II levels compared to control (**Fig. 4B**), albeit without any statistical significance (*p*=0.3426). However, in α-Syn–exposed neuronal cells treated with bafilomycin A1, LC3B-II levels were significantly lower than in cells treated with bafilomycin A1 alone, suggesting a reduction in autophagy flux. In microglial cells, by contrast, there was no significant reduction in LC3B-II levels upon α-Syn exposure relative to control (p = 0.9995), nor with co-treatment of α-Syn and bafilomycin A1 compared to bafilomycin A1 alone (p = 0.9501), suggesting an unperturbed flux (**Fig. 4C**).

**Figure 4.**
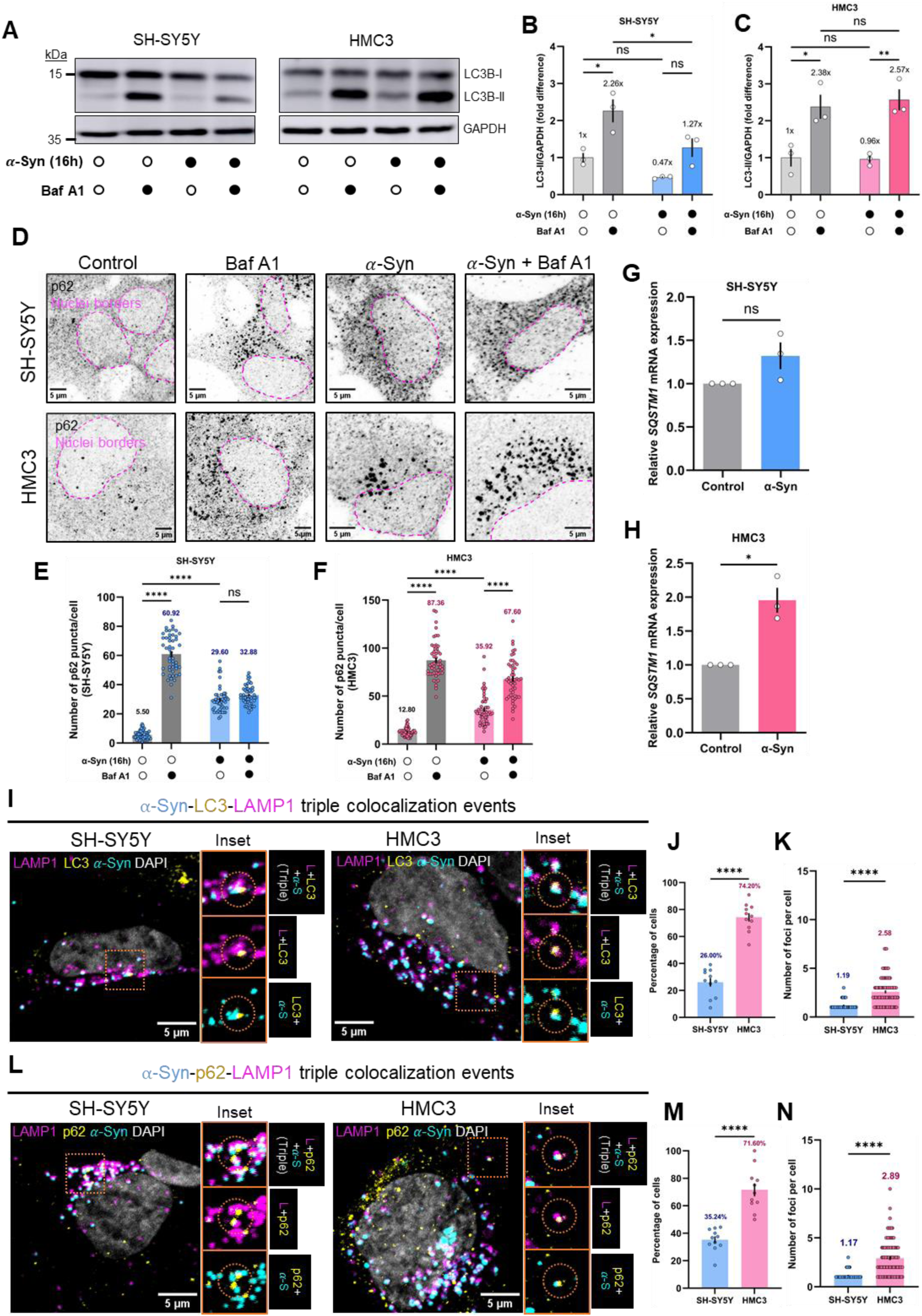
Effect of aggregates on autophagy flux, and its recognition as a cargo. (**A**) Representative immunoblot images of SH-SY5Y and HMC3 cells probed for LC3B. Filled circles represent treatment conditions. (**B and C**) Quantification of the relative level of LC3B-II in different conditions for SH-SY5Y (**B**) and HMC3 (**C**) cells, represented as fold difference of control. N=3 independent experiments. Error bars represent SEM. Statistical significance was analyzed using Two-Way ANOVA with Tukey’s multiple comparison. For (**B**): control versus Baf A1, *p=0.0118; control versus *α*-Syn, ns (p=0.3426); *α*-Syn versus *α*-Syn+Baf A1, ns (p=0.1022); and Baf A1 versus *α*-Syn+Baf A1, *p=0.0402. For (**C**): control versus Baf A1, *p=0.0197; control versus *α*-Syn, ns (p=0.9995); *α*-Syn versus *α*-Syn+Baf A1, **p=0.0085; and Baf A1 versus *α*-Syn+Baf A1, ns (p=0.9501). (**D**) Representative immunofluorescence images of p62 in the presence of *α*-Syn aggregates, with or without bafilomycin A1 (Baf A1) in SH-SY5Y cells (upper panels) and HMC3 cells (lower panels). Dotted lines demarcate borders of nuclei. (**E**) Corresponding quantification of the number of p62 puncta per cell for SH-SY5Y cells. (**F**) Corresponding quantification of the number of p62 puncta per cell for HMC3 cells. For both (**E**) and (**F**), N=3 independent experiments, n=50 cells per group. Error bars represent SEM. Mean values are mentioned within the graphs. Statistical significance was analyzed using Two-Way ANOVA with Tukey’s multiple comparison. ns: non-significant (p=0.2175), ****p<0.0001. (**G**) Relative transcript expression of SQSTM1 in SH-SY5Y cells. N=3 biological replicates. Error bars represent SEM. Statistical significance was analyzed using two-tailed Student’s t-test with Welch’s correction. ns: non-significant (p=0.1734). (**H**) Relative transcript expression of SQSTM1 in HMC3 cells. N=3 biological replicates. Error bars represent SEM. Statistical significance was analyzed using two-tailed Student’s t-test with Welch’s correction. *p=0.0355. (**I**) Representative images of neuronal (top panels) and microglial (bottom panels) cells depicting triple colocalization of *α*-Syn (cyan) with autophagy receptor LC3 (yellow) and lysosomes (magenta). Inset images to the right depict triple colocalization event (top panel), and dual-channel images of lysosome with LC3, or *α*-Syn. Orange dashed circle highlights the colocalized event. (**J**) Quantification of the percentage of cells per field-of-view with at-least one triple colocalization event (*α*-Syn with LC3 and LAMP1) in SH-SY5Y and HMC3 cells. Mean percentage is mentioned within the graph. N=3 independent experiments, number of cells analyzed: n=212 for SH-SY5Y and 133 for HMC3 from 11 independent images. (**K**) Average number of triple-colocalizing (aggregate with LC3 and LAMP1) foci per cell for SH-SY5Y and HMC3. Mean number of foci is mentioned within the graph. N=3 independent experiments, number of cells analyzed: n=56 for SH-SY5Y and 98 for HMC3 (cells positive for at least one triple colocalization event from [**J**]). (**L**) Same as in (**I**), but with p62 (yellow) as the autophagy receptor probed for. (**M**) Quantification of the percentage of cells per field-of-view with at-least one triple colocalization event (*α*-Syn with p62 and LAMP1) in SH-SY5Y and HMC3 cells. Mean percentage is mentioned within the graph. N=3 independent experiments, number of cells analyzed: n=201 for SH-SY5Y and 155 for HMC3 from 11 independent images. (**N**) Average number of triple-colocalizing (aggregate with p62 and LAMP1) foci per cell for SH-SY5Y and HMC3. Mean number of foci is mentioned within the graph. N=3 independent experiments, number of cells analyzed: n=69 for SH-SY5Y and 109 for HMC3 (cells positive for at least one triple colocalization event from [**M**]). Error bars represent SEM. Statistical significance was analyzed using unpaired Student’s t-test with Welch’s correction. ****p<0.0001.

The autophagic receptor p62/SQSTM1 (hereafter p62) has been shown to play a central role in aggregate clearance in different neurodegenerative diseases by sequestering cargos in condensates known as p62 bodies, eventually targeting them towards degradation (Kuusisto *et al*, 2001; Bjørkøy *et al*, 2005; Pankiv *et al*, 2007). Notably, under our experimental conditions, exposure to α-Syn in both neuronal cells and microglia for 16 hours led to a significant increase in the number of p62 puncta per cell (**Fig. 4D-F**). However, this augmentation could result from either an accumulation of p62 (indicative of autophagy flux dysfunction) or an upregulation of p62 expression (suggestive of autophagy flux upregulation). To discern between these possibilities, aggregate-loaded cells were treated with bafilomycin A1, or left untreated, followed by immunostaining for p62. In neuronal cells, no significant increase in the number of p62 puncta was observed upon bafilomycin A1 treatment in the presence of aggregates (**Fig. 4E**). In contrast, microglia cells showed a substantial increase in p62 puncta under the same conditions (**Fig. 4F**).

To further validate the status of autophagy flux upon *α*-Syn exposure, we assessed the expression of the autophagy target gene of TFEB, *SQSTM1* (**Fig. 4G, H**). Consistent with the increase in p62 puncta, we also observed an upregulation of *SQSTM1* expression in microglial cells upon aggregate exposure. Taken together, these results suggest that autophagy flux is impaired in neuronal cells upon *α*-Syn exposure, whereas it is upregulated in microglia.

Having observed a difference in autophagy flux between the two cell types in the presence of aggregates, we next investigated whether these aggregates are targeted to lysosomes. To this end, we co-immunostained for LC3 and LAMP1 (**Fig. 4I**) or p62 and LAMP1 (**Fig. 4L**) in neuronal cells (upper panels) and microglial cells (lower panels) exposed to *α*-Syn aggregates. We quantified the fraction of cells displaying triple localization events (aggregates co-localizing with an autophagy marker and lysosome), as well as the number of such events per cell. Both autophagy markers participated in triple localization events (*α*-Syn-LC3-LAMP1 and *α*-Syn-p62-LAMP1) in both neuronal cells and microglia. While over 70% of microglia exhibited triple localization events with both LC3 and p62, neuronal cells showed only 26% of triple localization with LC3, and 35.24% with p62 (**Fig. 4J and M**). In addition, the average number of triple-localized foci per cell was significantly higher in microglia compared to neuronal cells (**Fig. 4K and N**). In line with our previous observations, these results point towards a more efficient microglial autophagic pathway and associated targeting of *α*-Syn aggregates compared to neuronal cells.

### Human iPSC-derived neurons and microglia display similar responses to *α*-Syn aggregates as cell lines

To validate our findings in a more relevant model system, we derived control dopaminergic neurons (DAn) and microglia (MG) from induced pluripotent stem cells (hiPSC). DAn and microglia were generated following previously published protocols (Calatayud *et al*, 2019; Blasco-Agell *et al*, 2024). After 16 hours of exposure to *α*-Syn aggregates, a higher proportion of neuronal lysosomes localize with *α*-Syn aggregates compared to microglia (45.45% versus 13.05%) (**Fig. 5A-D**), consistent with our initial observations in cultured cell lines (**Fig. 1**).

**Figure 5.**
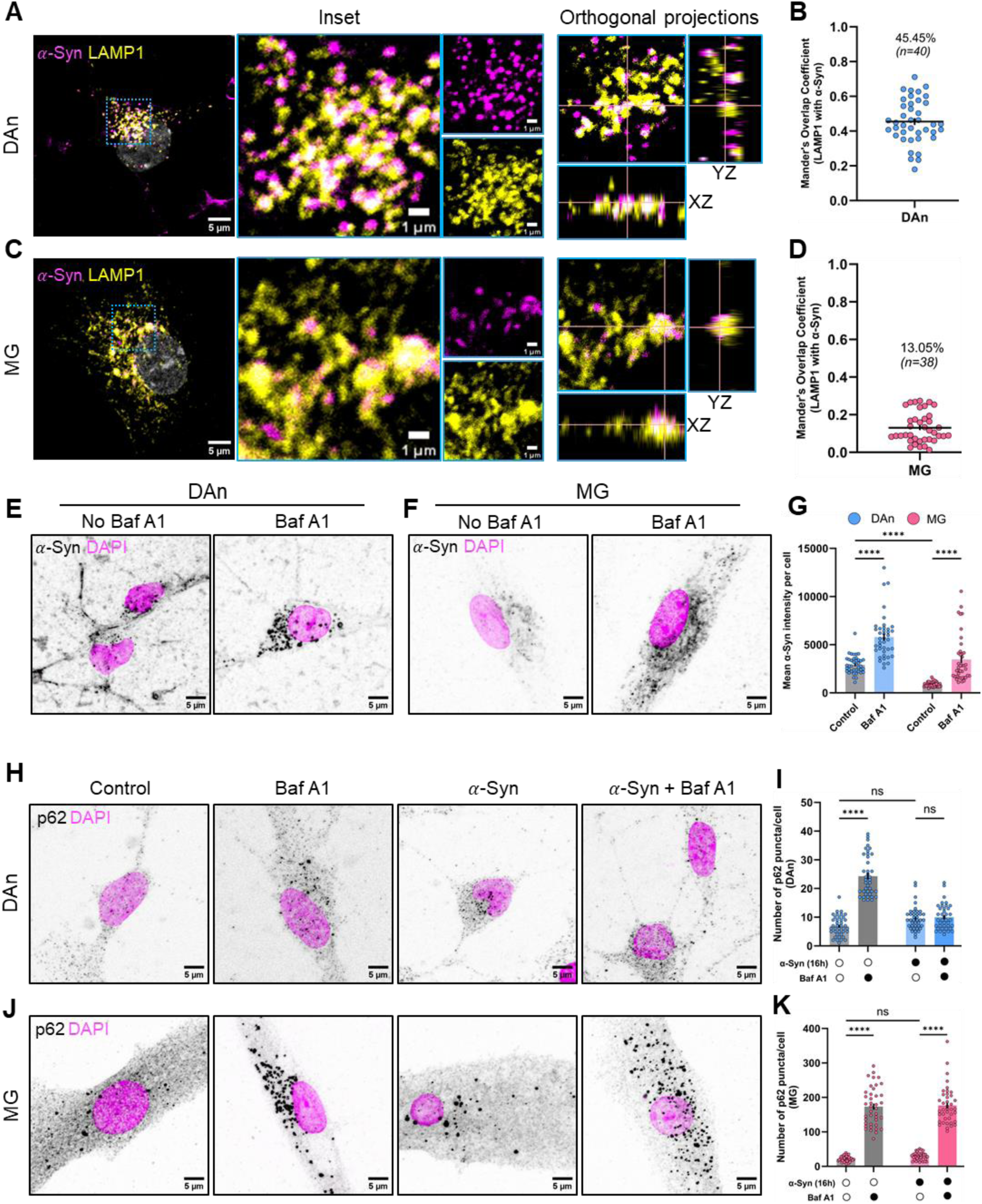
Validation of the neuronal and microglial responses to *α*-Syn using hiPSC-derived cells. (**A**) Representative images of *α*-Syn colocalization with LAMP1+ lysosomes in control neurons (DAn). Blue dashed box depicts region in the inset, orthogonal projection of which is shown. (**B**) Quantification of the percentage of lysosomes overlapping with *α*-Syn in control neurons. N=3 independent experiments, n=40 cells. Mean percentage mentioned within the graph. (**C**) Representative images of *α*-Syn colocalization with LAMP1+ lysosomes in control microglia. Blue dashed box depicts region in the inset, orthogonal projection of which is shown. (**D**) Quantification of the percentage of lysosomes overlapping with *α*-Syn in control microglia. N=3 independent experiments, n=38 cells. Mean percentage mentioned within the graph. (**E and F**) Representative images of *α*-Syn load in control neurons (**E**) and control microglia (**F**) in control and bafilomycin A1 treated conditions. (**G**) Quantification of the mean fluorescence intensity of *α*-Syn in different conditions. N=3 independent experiments, n=40 cells per group. Error bars represent SEM. Statistical significance was analyzed using Brown-Forsythe and Welch One-Way ANOVA with Dunnett’s T3 multiple comparison. ****p<0.0001. (**H**) Representative images of p62 immunostaining in control neurons in different conditions. (**I**) Quantification of the number of p62 puncta per cell in control neurons. N=3 independent experiments, n=40 cells per group. Error bars represent SEM. Statistical significance was analyzed using Brown-Forsythe and Welch One-Way ANOVA with Dunnett’s T3 multiple comparison. ns: non-significant, ****p<0.0001. (**J**) Representative images of p62 immunostaining in control microglia in different conditions. (**K**) Quantification of the number of p62 puncta per cell in control microglia. N=3 independent experiments, n=40 cells per group, except for only *α*-Syn treated group, for which n=38 cells. Error bars represent SEM. Statistical significance was analyzed using Brown-Forsythe and Welch One-Way ANOVA with Dunnett’s T3 multiple comparison. ns: non-significant, ****p<0.0001.

We attributed the lower percentage of *α*-Syn-LAMP1 colocalization in microglia to reduced number of aggregates in these cells, a factor that critically influences object-based colocalization analyses (i.e., a lower number of aggregates per cell at the time of detection results in a reduced colocalization index). To test this hypothesis, we analyzed the mean fluorescence intensity of *α*-Syn per cell in the presence or not of bafilomycin A1 (**Fig. 5E and F**). Not only did we detect a higher level of aggregates upon bafilomycin A1 treatment, suggestive of these aggregates being cargoes of autophagic degradation, we also detect lesser aggregates in microglia relative to neurons at a basal state (**Fig. 5G**), suggesting that microglia are relatively efficient in clearing these aggregates, an observation in line with our previous data.

Furthermore, we assessed the impact of *α*-Syn aggregates on autophagy flux in these cells. After 16 hours of aggregate exposure, we did not detect any significant increase in the number of p62 puncta per cell relative to control conditions in both neurons (**Fig. 5H and I**), and microglia (**Fig. 5J and K**). However, bafilomycin A1 treatment did not increase any further the number of p62 puncta in neurons, but did so in microglia. Taken together, these results suggest that *α*-Syn aggregates impair autophagy flux in neurons, but not in microglia, mirroring our previous observations in cell lines (**Fig. 4**).

### Inhibited autophagy contributes to increased aggregate transfer

Building upon these data, as well as a prior study demonstrating a directional bias in the transfer of α-Syn aggregates from neuronal cells to microglia (Chakraborty *et al*, 2023b), we hypothesized that autophagy impairment in neuronal cells could promote the dissipation of aggregates from neuronal cells to microglia. To test this hypothesis, we utilized the irreversible autophagy inhibitor, wortmannin, which selectively inhibits phosphoinositide-3-kinase (PI3K) and prevents autophagosome biogenesis (Blommaart *et al*, 1997). Neuronal cells loaded with *α*-Syn aggregates were treated with wortmannin for 2 hours and then fixed either immediately or after an additional 12 hours of growth in drug-free media. We observed an increased load of aggregates within 2h of wortmannin treatment (1.27-folds higher), which further rose to 1.42-folds after 12h (**Fig. S3A, left panels and B**).

Having observed an increased aggregate accumulation in neuronal cells upon autophagy inhibition, we performed co-cultures of aggregate containing SH-SY5Y cells (N) pre-treated or not with wortmannin (Wm) and HMC3 cells (M) for 12h – (N-M versus N^Wm^-M) and assessed for *α*-Syn transfer from donor neuronal cells to acceptor microglia (**Fig. S4A, and 5A**). A higher percentage of acceptor microglia contained aggregates upon wortmannin treatment (N^Wm^-M) compared to no drug treatment (control, N-M) (**Fig. 6B**). As expected, autophagy inhibition in donor neuronal cells increased the extent of transfer to microglia (N^Wm^-M) by 29.07% compared to control. Additionally, we performed similar experiments by treating only the acceptor cells with wortmannin (N-M^Wm^), or both donor and acceptor cells (N^Wm^-M^Wm^) (**Fig. S4A, B**).

**Figure 6.**
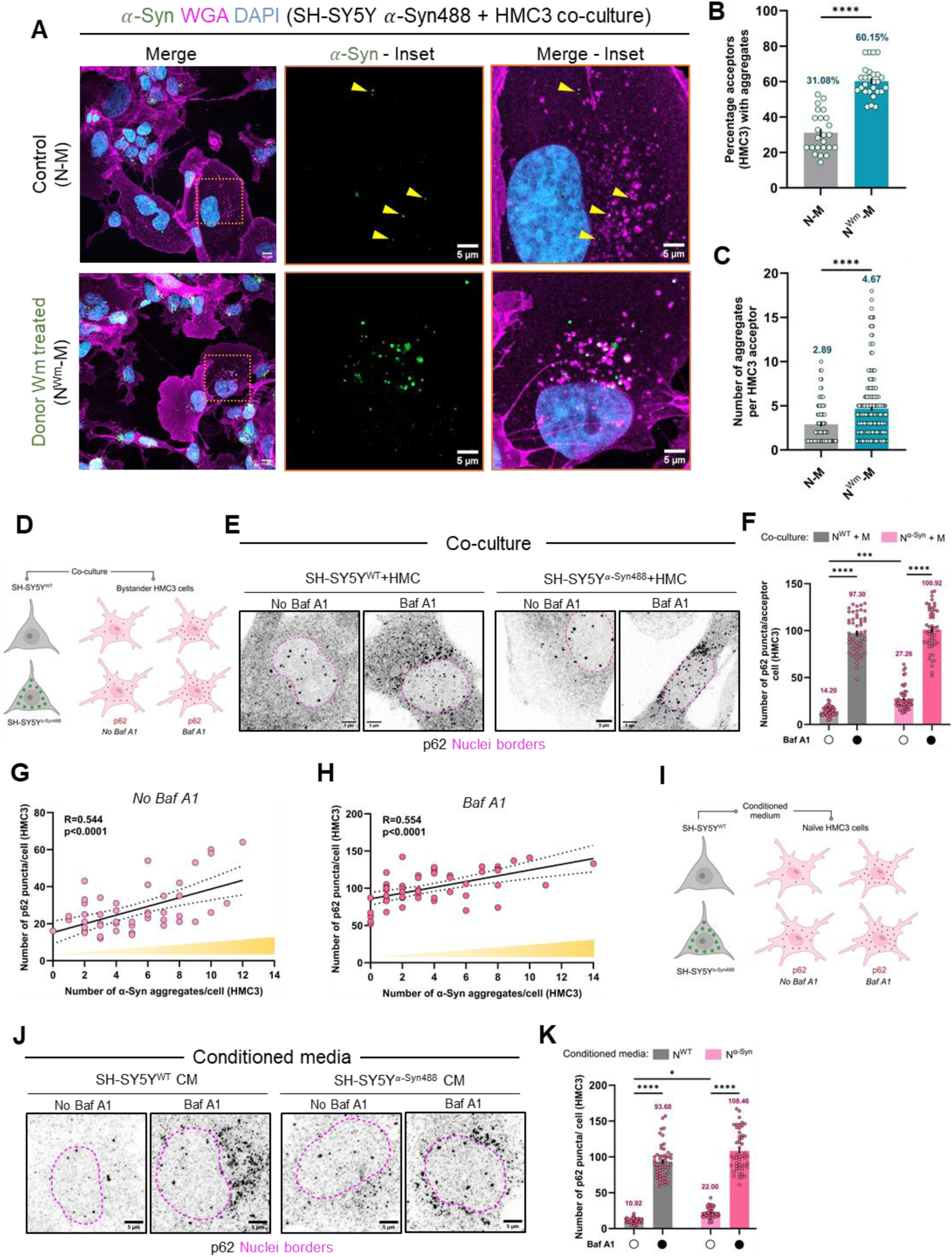
Contribution of autophagy in transfer of aggregates, and bystander influence. (**A**) Representative images of 12h co-cultures between *α*-Syn loaded SH-SY5Y neuronal cells and HMC3 microglia (control and wortmannin treated – see also figure EV4A for schema of the experimental set-up). Wortmannin treatment represented by the superscript “Wm”. Dotted box indicates the region of acceptor microglia zoomed in. (**B**) Percentage of HMC3 acceptor cells that received aggregates in control and wortmannin-treated cocultures. Average percentage of acceptor cells positive for aggregates is mentioned within the graph. N=3 independent experiments, n=240 acceptor cells for N-M, 240 acceptor cells for N^Wm^-M. Error bars represent SEM. Statistical significance was analyzed using Mann-Whitney U test. ****p<0.0001 (**C**) Quantification of the number of aggregates received per cell by acceptor HMC3 in (**B**). Mean number of aggregates is mentioned within the graph. N=3 independent experiments. N=137 acceptor cells for N-M, 196 acceptor cells for N^Wm^-M. Error bars represent SEM. Statistical significance was analyzed using Mann-Whitney U test. ****p<0.0001. (**D**) Schema depicts the co-culture condition of neuronal cells loaded or not with *α*-Syn488 aggregates and microglia, followed by immunostaining against p62 in basal or bafilomycin A1 treated conditions. (**E**) Representative p62 immunofluorescence (gray-invert LUT) images of HMC3 acceptor cells. (**F**) Quantification of the number of p62 puncta per microglial acceptor cell in co-culture conditions. N represents neuronal cells and M microglia. Mean values are mentioned within the graph. N=3 independent experiments, n=50 cells per group. Error bars represent SEM. Statistical significance was analyzed using Two-Way ANOVA with Šídák’s multiple comparison. ***p=0.0007, ****p<0.0001. (**G and H**) Correlation plot of the number of p62 puncta detected per microglial acceptor cells as a function of the number of *α*-Syn aggregates it received in basal (no baf A1, **G**) or autophagy inhibited (with baf A1, **H**) conditions. p-value represents he significance level of a non-zero slope, analyzed by simple linear regression. (**I**) Schema depicts the conditioned media experiment to assess secretory influence of neuronal cells on microglial autophagy, followed by immunostaining against p62 in basal or bafilomycin A1 treated conditions. (**J**) Representative p62 immunofluorescence (gray-invert LUT) images of HMC3 cells. (**K**) Quantification of the number of p62 puncta per microglial cell exposed to different conditioned media. N represents neuronal cells and M microglia. Mean values are mentioned within the graph. N=3 independent experiments, n=50 cells per group. Error bars represent SEM. Statistical significance was analyzed using Two-Way ANOVA with Šídák’s multiple comparison. *p=0.0207, ****p<0.0001.

Interestingly, a comparable increase in aggregate transfer was observed in both experimental conditions of wortmannin treatment (**Fig. S4C**). This could be due to either increased transfer, and/or decreased clearance of aggregates in the treated microglia. To distinguish between these possibilities, we measured the aggregate load in microglia either immediately after 2h of wortmannin treatment, or after 12h of culture in drug-free media. We found an elevated load of aggregates in microglia following autophagy inhibition (**Fig. S3A, right panels and C**), consistent with the essential role of autophagy in the clearance of *α*-Syn aggregates. Together with the observed increase in aggregate transfer, the higher levels of *α*-Syn in the N-M^Wm^ group could therefore be attributed to impaired clearance of aggregates by microglia after receiving them. Importantly, wortmannin treatment did not result in increased exocytosis of aggregates, and the extent of aggregate transfer via secretion was much lower than that observed in co-cultures (**Fig. S3D**).

Upon autophagy inhibition, not only were more acceptor cells positive for aggregates, but there was also an increase in the average number of aggregates per acceptor cell (**Fig. 6C**). In the control condition, the average number of aggregates per acceptor microglia was 2.89, whereas upon wortmannin treatment of donor cells (N^Wm^-M) microglia contained 4.67 aggregates. When autophagy was inhibited in the acceptor microglia (N-M^Wm^), this number increased further to 6.27 aggregates per cell **(Fig. S4D)**. Interestingly, wortmannin treatment of both donor and acceptor cell populations (N^Wm^-M^Wm^) resulted in the highest number of aggregates present in acceptors, with an average of 11.84 per acceptor cell (**Fig. S4D)**. This suggests an additive effect on the quantity of aggregates transferred when both populations are treated with wortmannin. Together, these results support the idea that autophagy dysfunction plays a causal role in the transfer of aggregates from neuronal cells to microglia.

As a proof-of-concept that autophagy impairment leads to increased aggregate transfer, we performed a similar experiment, but with microglia serving as the donor cells (**Fig. S5A**). Consistent with our previous results (Chakraborty *et al*, 2023b), in control conditions we observed less transfer of aggregates from microglia to neuronal cells (4.80% in M-N group), which increased to 13.34% upon autophagy impairment (M^Wm^-N) (**Fig. S5B and C**).

As we did not observe any change in the extent of aggregate release via secretion upon wortmannin treatment (**Fig. S3D**), we next assessed if autophagy dysfunction is associated with an increase in TNT connections between cells. After treating both neuronal and microglial cells with wortmannin, we found an increase of 1.83-folds for SH-SY5Y (**Fig. S6A and B**) and of 1.67-folds for HMC3 cells (**Fig. S6C and D**) in the extent of connections. To further validate whether autophagy impairment increases intercellular connections, we used bafilomycin A1 which inhibits autophagy at a later step— autophagosome-lysosome fusion—in contrast to wortmannin, which inhibits autophagy initiation. Upon bafilomycin A1 treatment of cells, the number of connections increased by over 1.5-fold for both cell types (**Fig. S6E-H**). Taken together, these data suggest that autophagy impairment plays a functional role in regulating intercellular connections, a major route for *α*-Syn transfer between neuronal and microglial cells.

### Neuronal cells with *α*-Syn induce autophagy in bystander microglia

Cells grown in co-cultures often behave differently compared to those grown in monocultures. The phenomenon of bystander influence on autophagy has been observed during irradiation, known as radiation-induced bystander effect (RIBE) (Huang *et al*, 2014; Song *et al*, 2016). Additionally, chemokines secreted by activated microglia have been reported to inhibit neuronal autophagy in neurodegenerative pathologies (Festa *et al*, 2023). To investigate potential non-cell autonomous effects in our system, we co-cultured microglia with neuronal cells loaded or not with *α*-Syn aggregates, and performed immunostaining against p62. The flux was determined by treating co-cultures with bafilomycin A1 (**Fig. 6D and E**). We observed a significant increase in the number of p62 puncta per microglial cell when cultured with aggregate-burdened neuronal cells (N*α*^-Syn^ + M; average of 27.26), compared to co-cultures with healthy neuronal cells (N^WT^ + M; average of 14.20) (**Fig. 6F**). A larger increase in microglial p62 accumulation was observed upon bafilomycin A1 treatment, indicating upregulated autophagy flux. Interestingly, we found a positive correlation between the number of p62 puncta per microglial cell and the number of *α*-Syn aggregates it received from neuronal cells (**Fig. 6G and H**). To assess whether such a phenotype was only contact-mediated, we performed a similar experiment on monocultures of microglial cells exposed to conditioned medium from naïve or *α*-Syn-laden neuronal cells (**Fig. 6I**). We observed a similar increase in the number of p62 puncta per cell when exposed to conditioned media of aggregate-exposed neuronal cells (N*α*^-Syn^; average of 22.00) relative to that of healthy cells (N^WT^; average of 10.92) (**Fig. 6J and K**). Taken together, these data suggest the influence of *α*-Syn-burdened neuronal cells on autophagy flux of microglial cells via both contact-dependent and independent mechanisms.

To assess the relevance of this upregulated autophagy in microglia, we tracked the fate of aggregates post-transfer, and observed 38.57% of transferred aggregates co-localized with p62 in microglial cells (**Fig. 7A-C**), suggesting that neuronal cell-derived aggregates are destined for degradation in microglia.

**Figure 7.**
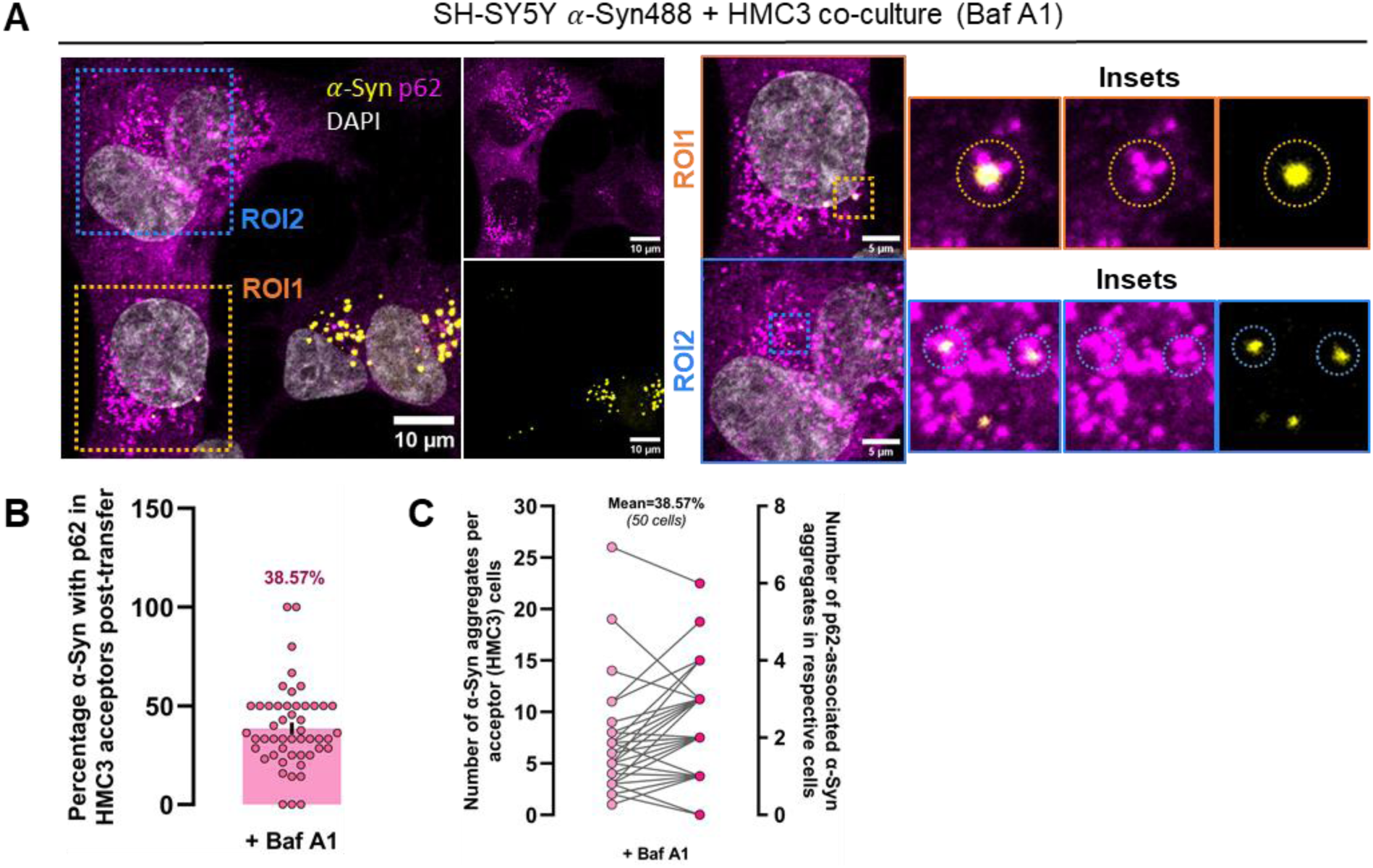
Fate of *α*-Syn aggregates post-transfer. (**A**) Representative images of *α*-Syn-loaded neuronal cells in co-culture with acceptor microglia immunostained for p62 in baf A1-treated condition. ROIs 1 and 2 are zoomed for regions of *α*-Syn colocalizing with p62 (insets). (**B**) Quantification of the percentage of *α*-Syn colocalized with p62 in microglia post-transfer from neuronal cells. Mean value is mentioned within the graph. (**C**) Similar quantification as in (**G**), but with details of the number of *α*-Syn recognized by p62 per acceptor cells (right y-axis) and the number of total aggregates received (left y-axis). For both **(B)** and (**C**), N=3 independent experiments, n=50 acceptor cells analyzed.

## Discussion

### Role of TNTs in aggregate spreading

Several studies have reported that amyloidogenic aggregates can induce the formation of intercellular connections such as tunneling nanotubes (TNTs), which facilitate their spread between cells. Initially, this was demonstrated with aggregated prion protein (PrPSc) between CAD neuronal cells and dendritic cells (Gousset *et al*, 2009). More recently, research has found that various aggregates, including Amyloid-β, Tau, α-Syn, and mutant Huntingtin (mHTT), increase intercellular connections, further promoting aggregate transfer (Chakraborty *et al*, 2023a). Notably, TNT-like connections are increased *in vitro* by the small GTPase, Rhes (Sharma & Subramaniam, 2019). Given its enrichment in the striatum, Rhes also facilitates mHTT spread *in vivo* from the striatum to cortical areas (Ramírez-Jarquín *et al*, 2022).

*In vitro, α*-Syn aggregates are transferred more efficiently from mouse neurons to astrocytes than in the opposite direction (Loria *et al*, 2017). Such transfer has also been demonstrated between murine primary cortical neurons and microglia (Scheiblich *et al*, 2024). Similarly, human neuronal cells transfer *α*-Syn aggregates to microglia more effectively via TNTs than they receive them (Chakraborty *et al*, 2023b). These studies suggest a conserved mechanism allowing neurons to offload their aggregate burden via TNTs.

Here, we uncover a causative role of autophagy dysfunction in driving TNT-mediated *α*-Syn aggregate transfer from neuronal cells to microglia. This provides a mechanistic insight into the directionality of aggregate transfer, while revealing differential impacts of α-Syn aggregates on the lysosomal and autophagic pathways in neuronal versus microglial cells.

### Differential effect of *α*-Syn aggregates on lysosome function and autophagic pathways in neuronal cells versus microglia

Impaired autophagy is a molecular hallmark of neurodegenerative diseases (Wilson *et al*, 2023). The clearance of aggregate-prone proteins, such as *α*-Syn, heavily relies on autophagy, although the ubiquitin-proteasome system also play a role (Webb *et al*, 2003; Vogiatzi *et al*, 2008). Notably, inhibition of autophagosome-lysosome fusion using bafilomycin A1 increases *α*-Syn-associated cytotoxicity (Klucken *et al*, 2012). This sensitivity to bafilomycin A1 correlates with reduced levels of SNAP29, a SNARE protein essential for autophagosome-lysosome fusion, in α-Syn-transduced cells (Tang *et al*, 2021).

*In vivo,* in MPTP models of PD, the loss of dopaminergic neurons is preceded by lysosomal dysfunction and a reduced lysosome number, leading to autophagosome accumulation. These phenotypes are rescued by rapamycin-induced autophagy activation (Dehay *et al*, 2010). Similarly, decreased lysosome-associated markers have been reported in human PD subjects (Chu *et al*, 2009). Further studies have shown impaired lysosomal functions in neuronal cells treated with exogenous *α*-Syn aggregates and in PD patient iPSC-derived dopaminergic neurons (Senol *et al*, 2021; Drobny *et al*, 2023). Additionally, lysosomes containing *α*-Syn are shuttled between cells through TNTs (Senol *et al*, 2021; Abounit *et al*, 2016). Based on these findings, we hypothesized that impaired autophagy in *α*-Syn-treated neuronal cells contributes to the spread of aggregates to neighboring cells via TNTs.

Our results demonstrate that α-Syn induces greater lysosomal damage in neurons than in microglia, as evidenced by increased aggregate association with lysosomes **(Fig. 1B, C)** and reduced neuronal lysosomal degradation capacity, which, on the contrary, is retained for microglia **(Fig. 2).** Similar to what happens in cell lines the addition of α-syn PFFs also induces differential localization of *α*-Syn to lysosomes in hiPSC-derived neurons and microglia (**Fig. 5A-D**) highlighting a conserved biological process between cell lines and near-physiological iPSC-derived cells. The heightened functionality of microglial lysosomes in the presence of aggregates is likely due (or consistent) to microglia’s role as brain macrophages. However, microglial lysosomes are less acidic than those of macrophages (Majumdar *et al*, 2007; Quick *et al*, 2023), limiting their degradative capacity. Acidity and functionality of these organelles increase in pro-inflammatory environments, as observed in fibrillar Amyloid-β clearance (Majumdar *et al*, 2007) and TLR2-mediated autophagy activation in microglia (Arroyo *et al*, 2013). Thus, the enhanced degradative ability of microglial lysosomes in our experiments may result from an α-Syn-induced pro-inflammatory environment. However, further investigations are needed to confirm this hypothesis.

Additionally, the observed phenotype in microglia could be attributed to elevated lysosome biogenesis and/or lysosome repair pathways, which lead to a homeostatic maintenance/upregulation of lysosome functionality. Previous research on BV2 microglia demonstrated that upon treatment with fibrillar *α*-Syn, TANK-Binding Kinase 1 (TBK1) and Optineurin (OPTN) are recruited on lysosomes, facilitating the autophagic clearance of damaged organelles (Bussi *et al*, 2018). Consistent with these findings, we observe elevated lysophagy and concurrent lysosome biogenesis in microglia compared to neuronal cells (**Fig. 3**). This dynamic turnover of lysosomes suggests a mechanism by which microglia are able to counteract the detrimental effects of *α*-Syn aggregates on lysosomal function.

Given these differences, we hypothesize a differential autophagy response to *α*-Syn aggregates between the two cell types. We observed upregulated autophagy in both microglia cell line and hiPSC-derived microglia (**Fig. 3A, C; and Fig. 5J, K**), as opposed to dysfunctional autophagy and accumulation of autophagy substrate p62 in neuronal cell lines and hiPSC-derived neurons (**Fig. 4A-H and Fig. 5E-K**). Enhanced aggregate co-localization with autophagy markers LC3 and p62, as well as with LAMP1 in microglia corroborates previous studies showing upregulated autophagy in BV2 microglia exposed to fibrillar α-Syn (Bussi *et al*, 2018). These findings underscore critical differences in the cellular responses to α-Syn aggregates, indicating that neurons are unable to clear aggregates as efficiently as microglia. This deficit in neuronal autophagy may contribute to the impaired clearance of α-Syn aggregates, further exacerbating neurodegenerative processes in conditions such as Parkinson’s disease.

These findings align with a recent study showing that microglia efficiently clear neuron-released α-Syn aggregates in a p62-dependent manner (Choi *et al*, 2020). The authors also identified a functional link between the toll-like receptor 4 (TLR4)-nuclear factor kappa-light-chain-enhancer of activated B cells (NF-*κ*B) pathway and autophagy-dependent aggregate clearance. NF-*κ*B, a key transcription factor for pro-inflammatory cytokines production during *α*-Syn pre-formed fibrils exposure (Dutta *et al*, 2021), also regulates autophagy by inducing the expression of related genes (Nivon *et al*, 2009; Copetti *et al*, 2009). Thus, the observed upregulation of autophagy in microglia might result from NF-*κ*B activation through toll-like receptors, although this requires further investigation. Taken together, these results highlight the need for neuronal cells to find a route to dissipate the burden of aggregates.

### Autophagy Dysregulation Drives Aggregate Transfer

Given the significant impact of aggregate accumulation on autophagy in neuronal cells, we hypothesized that this impairment could contribute to the transfer of aggregates to microglial cells (Chakraborty *et al*, 2023b). Inhibition of autophagy in aggregate-laden neuronal cells by wortmannin resulted in a notable increase in aggregate load (**Fig. S3A-C**), and transfer to microglia (**Fig. 6A-C**) underscoring the role of dysfunctional autophagy in the dissipation of aggregates to neighboring cells. Conversely, wortmannin treatment of acceptor microglia (N-M^Wm^), or both donors and acceptors (N^Wm^-M^Wm^) resulted in an increase in α-Syn-aggregates compared to the control, likely reflecting the cumulative effect of elevated transfer and accumulation of aggregates not targeted for degradation (**Fig. S4B and C**). Interestingly, the number of aggregates transferred increased when both donors and acceptors were treated (**Fig. S4D**). Assuming a linear relationship between autophagy inhibition and aggregate transfer, a predictive value for the average number of aggregates transferred (calculated by adding the effects of wortmannin on individual cell populations) was 10.94 (N^Wm^-M + N-M^Wm^ groups), closely matching the observed value of 11.84 (N^Wm^-M^Wm^ group). Nevertheless, it is yet to be determined whether there is bi-directional communication between neuronal cells and microglia that influences the extent of contact-mediated aggregate transfer upon autophagy inhibition. Of interest, a recent study highlights a non-cell autonomous mechanism in Huntington’s disease (HD) and tauopathies, where activated microglia secrete proinflammatory chemokines (CCL-3/-4/-5) that suppress neuronal autophagy via mTORC1 activation through CCR5 (Festa *et al*, 2023). Our findings suggest that aggregate-laden neurons with impaired autophagic flux induce a compensatory autophagic response in microglia. Microglia cultured with aggregate-loaded neurons displayed enhanced autophagy, correlating with increased aggregate uptake **(Fig. 6D-H)**. This phenotype was also observed when microglial cells were exposed to neuronal conditioned media (**Fig. 6I-K**). These results highlight the critical role of bystander effects in the cellular response to *α*-Syn aggregates, particularly within microglia, suggesting a non-cell autonomous mechanism of aggregate clearance. The upregulation of autophagy in microglia, triggered by aggregate transfer, suggests that microglia play a key role in clearing neuronal-derived pathological aggregates, potentially restoring proteostasis in the surrounding environment. The precise influences of contact-mediated interactions or secretory factors in eliciting a robust non-cell autonomous autophagy response in microglial cells remain to be determined.

Additionally, aggregate transfer from microglia to neurons also increased upon autophagy inhibition **(S5B-D).** Although transfer from microglia to neurons is typically less efficient, this finding supports the hypothesis that autophagy dysfunction serves as a signaling mechanism for intercellular aggregate transfer. Furthermore, autophagy inhibition using wortmannin and bafilomycin A1 significantly elevated intercellular connections between neuronal and microglial cells **(Fig. 5D-F and Fig. S5E-H).** This increase can be attributed to the accumulation of superfluous materials caused by autophagy inhibition, which triggers a global stress response promoting TNT formation (Kroemer *et al*, 2010).

Mechanistically, autophagy is closely linked to the endoplasmic reticulum (ER) stress response. While ER stress typically induces autophagy, this regulation is disrupted under several pathological conditions, including neurodegenerative diseases (Rashid et al., 2015). ER stress activation leads to cytosolic Ca²⁺ release via the inositol trisphosphate (IP3) receptor. Elevated cytosolic Ca²⁺ levels have been shown to positively regulate TNT formation. In particular, Ca²⁺/calmodulin-dependent protein kinase II (CaMKII), a key transducer of the Wnt/Ca²⁺ pathway, regulates TNT formation in neuronal cells through the actin-binding activity of its β-isoform (Vargas et al., 2019).

Additionally, impairment of autophagosome–lysosome fusion has been reported to induce lysosomal membrane permeabilization (LMP) (Hu et al., 2021)—a phenomenon we observed to be significantly more prominent in neuronal cells than in microglia (**Fig. S1A, C, D)**, leading to autophagosome accumulation. Interestingly, this accumulation was reversed by treatment with BAPTA-AM, a calcium chelator, suggesting a role for Ca²⁺ in this process.

Although the precise mechanism of TNT formation remains incompletely understood, actin polymerization into long, parallel filaments is a critical early step. These filaments allow TNTs to span distances beyond those of typical cellular protrusions, forming stable intercellular connections (Henderson et al., 2023). This actin-driven architecture facilitates the transfer of cargo such as α-Syn aggregates and is also regulated by the Rac–PAK signaling pathway, which may be activated by aggregates (Scheiblich et al., 2024).

Together, our findings support a mechanistic link between autophagy dysfunction, ER stress, and TNT formation—a regulatory network that remains largely unexplored and may be central to the intercellular spread of neurodegenerative pathology.

## Conclusion and Future Directions

Our study reveals a mechanism underlying elevated aggregate transfer between cells, emphasizing the role of autophagy dysfunction in the propagation of aggregate-prone proteins. It also lays the foundation for further exploration of lysosome-regulated TNT-mediated communication between cells, in both physiological and pathological contexts. These findings underscore the importance of intercellular communication and autophagic flux modulation in the pathogenesis of neurodegenerative diseases, where impaired degradation of *α*-Syn in neurons drives aggregate transfer to microglia, which attempt to mitigate neuronal damage through enhanced autophagic activity. Targeting autophagy pathways may offer therapeutic potential to not only reduce the burden of α-Syn aggregates at an intra-cellular level, but also limit the inter-cellular spread of toxic aggregates, thereby mitigating the progression of the disease. Importantly, our observations may extend beyond interactions between neuronal and microglial cells to include neuron-astrocyte communication, offering a more comprehensive understanding of the biological underpinnings of these diseases.

## Limitations of the study

Due to the inherent difficulty of transfecting microglial cells for overexpression or knockdown studies, we conducted all experiments without genetic perturbations, relying instead on relevant and well-characterized pharmacological agents. While we validated our findings in iPSC-derived neurons and microglia, future studies employing advanced 3D cellular models (such as brain organoids and assembloids) as well as complementary animal models carrying patient-specific or autophagy-related gene mutations, will be essential to confirm these results in more complex systems and to further assess their relevance to human disease pathogenesis.

## Materials and Methods

### Culture of Cell lines

Human neuroblastoma cell line SH-SY5Y (referred to as neuronal cells in the manuscript, derived from SK-N-SH neuroblastoma line) were cultured in RPMI1640 media (Euroclone, ECB2000L), and supplemented with 10% foetal calf serum (FCS) (Eurobio Scientific, CVFSVF00-01) and 1% penicillin-streptomycin (Gibco, 15140-122; 100 units/mL final concentration). The human microglia clone 3 cell line (HMC3) were cultured in DMEM (Sigma-Aldrich, D6429) supplemented with 10% FCS and 1% penicillin-streptomycin. Cells were cultured in a humidified CO_2_ incubator at 37°C, and passaged using 0.05% Trypsin-EDTA (Gibco, 25300-054) when they reached 90% confluency. Cells were counted before each experiment, and seeded on autoclaved, 12 mm glass coverslips (uncoated) (Epredia, CB00120RA120MNZ0) for fixed-cell imaging, and on 35 mm glass-bottom microdishes (Ibidi GmbH, 81156) for live-cell imaging. Cells between passages 3 and 10 were used for experiments.

### Cell culture experiments with hiPSC lines

All procedures adhered to Spanish and EU guidelines and regulations for research involving the use of human pluripotent cell lines. The human iPSC lines used in our studies were generated following procedures approved by the Commission on Guarantees concerning the Donation and Use of Human Tissues and Cells of the Carlos III Health Institute, Madrid, Spain.

For both neurons and microglia, control hiPSC lines were used (SP11 and SP13wt/wt, respectively). The hiPSC line generation and characterization is as previously described (Sánchez-Danés *et al*, 2012; di Domenico *et al*, 2019). hiPSC were maintained on Matrigel (Corning, 354234)-coated plastic plates (Thermo Fisher Scientific) and in mTeSR^TM^1 medium (Stem Cell Technology, 85850) until the start of the protocols of differentiation.

The employed protocol for the differentiation of microglia and its characterization are described in detail in (Blasco-Agell *et al*, 2024). Briefly, hiPSC were passed in single colonies and, after 2-4 days, mTeSR^TM^1 medium was supplemented with 80 ng/mL of Bone Morphogenetic Protein (BMP)-4 (PeproTech, 120-05) for a total of 4 days. From the following day, cells were changed using SP34 medium (StemProTM-34 SFM (Gibco™, 10639011), 1% of P/S (Cultek, SV30010) and 1% of Ultraglutamine (Glut; Lonza, LZBE17-605EU1). For two days, media was supplemented with 80 ng/ml of VEGF (PeproTech, AF-100-20), 100 ng/ml of Stem Cell Factor (SCF, PeproTech, 300-07) and 25 ng/ml of Fibroblast Growth Factor (FGF)-2 (PeproTech, 100-18B). Then, from day 7 to day 14, supplemented factors included 50 ng/ml of Fms-like tyrosine kinase 3-Ligand (Flt3-L, Humanzyme, HZ-1151), 50 ng/ml of IL-3 (PeproTech, 200-03), 50 ng/ml of SCF, 5 ng/ml of Trombopoietin (TPO, PeproTech, 300-18) and 50 ng/ml of M-CSF (PeproTech, 300-25). The last step consisted on the addition to SP34 medium of Flt3-L, M-CSF and 25 ng/ml of GM-CSF (PeproTech, 300-03), changing the medium every 3-4 days. Starting from day 35, floating microglial progenitors were collected from the culture’s supernatant and passed through a 70 μm Filcon™ Syringe-Type nylon mesh (BD Biosciences, 10271120). Cells were counted and centrifuged at 300x*g* for 10 minutes. Recollected progenitor microglial cells were plated at a final density of 5.000 microglia in a well of a 24 well plate (Thermo Fisher Scientific) and on top of plastic coverslips. Media was changed twice a week with Roswell Park Memorial Institute (RPMI) 1640 Medium (Gibco^TM^, 11875093) supplemented with 50 ng/mL of IL-34 (PeproTech,200-34) and M-CSF. Microglia were considered mature after one week in culture.

NPCs were generated following a previously published protocol from (Tristán-Noguero *et al*, 2023; Chambers *et al*, 2009). Briefly, iPSCs were split into a 96 well-plate, V-bottom shape and centrifuged 800xg for 10 minutes to force the aggregation. Cells were grown on mTeSR medium (STEMCELL technologies, 05825) for 24 hours. Embryoid bodies (EBs) were plated in a 6 cm^2^ dish and the medium was then changed to Proneural [DMEM/F12 (Life, 21331-20) and Neurobasal (Life, 21103-049) – 1:1, 0.5% N2 (Life, 17502048), 1% B27 w/o Vitamin A (Life, 12587-010), 1% L-Glutamine (Linus, X0551-100), 1% Penicillin/Streptomycin (ScienCell, 0503), and 2-Mercaptoethanol (gibco 31350-010)]. EBs were seeded in POLAM-coated (poly-L-ornitine Sigma-aldrich, P4957; laminin, Sigma-Aldrich, L2020) wells of a 6 well plate with Proneural supplemented with Noggin 200ng/ml (PeproTech, 120-10C) and SB431542 10 µM (TOCRIS, 301836-41-9). When NEP rosettes were visible, they were enzymatically dissociated with trypsin 0.05% to obtain NPCs and plated on POLAM-coated wells of a 12 well plate. NPCs were then split up to 6-8 times in order to purify the culture.

DAn neurons were differentiated following a previously published protocol (Calatayud *et al*, 2019). NPCs at 80-100% confluency were cultured on POLAM-coated (poly-L-ornithine Sigma-aldrich, P4957; laminin, Sigma-Aldrich, L2020) wells in DAn induction medium [DMEM/F12 (Life, 21331-20), 1% N2 (Life, 17502048), 1% Penicillin/Streptomycin (ScienCell, 0503)] supplemented with 200 ng/ml Sonic Hedgehog (PeproTech, 100-45) and 100 ng/ml FGF8 (PeproTech, 100-25) for 6 days. This step allowed for NPCs patterning towards dopaminergic fate. DAn progenitors were then plated on POLAM-coated dishes in N2B27 medium (DMEM/F12 (Life, 21331-20) – Neurobasal (Life, 21103-049) 1:1, 0.5% N2 (Life, 17502048), 1% B27(Life, 17504-044), 1% L-Glutamine (Linus, X0551-100), and 1% Penicillin/Streptomycin (ScienCell, 0503) for 10 days for maturation. For terminal differentiation DAn were cultured on Matrigel (Corning, 354234) coated wells supplemented with 20 ng/ml BDNF (Peprotech, 450-02) and 20 ng/ml GDNF (Peprotech, 450-10) for 25 days.

### Preparation of *α*-Syn aggregates

*α*-Syn aggregates were prepared as described before (Nonaka *et al*, 2010). Briefly, human wild-type *α*-Syn was purified from *Escherichia coli* BL21 (DE3) with RP-HPLC. Fibrils were then conjugated with Alexa Fluor™ 488 or 568 fluorophores (used in imaging experiments) (Invitrogen) using manufacturer’s labelling kit, or not tagged with a fluorophore. Fibrils were kept at -80°C for long-term storage. Prior to exposure to cells, fibrils were diluted in growth medium at a working concentration of 500 nM and sonicated (BioBlock Scientific, Vibra Cell 7504) for 5 minutes at an amplitude of 80%, pulsed for 5 seconds “on” and 2 seconds “off”. For iPSC-derived cells, owing to their fragile nature, a lower concentration of fibrils (200 nM) was used. Sonicated fibrils were then added on cells directly without the addition of any intracellular delivery agents for designated time points. Post-incubation, cells were washed with diluted trypsin solution (1:3, in 1X DPBS [Gibco, 14040-091]) and processed for experiments.

### Immunocytochemistry

Immunofluorescence on cells were performed using the standard protocol. Cells were fixed with 4% paraformaldehyde (PFA [Electron Microscopy Sciences, 15710]) for 30 minutes at room temperature (RT), followed by one wash with 1X DPBS. Cells were then incubated in 50 mM ammonium chloride (NH_4_Cl [Sigma Aldrich, A0171]) solution for 15 minutes at RT to quench the fixative. Following three washes with 1X DPBS, cells were incubated in 1X DPBS-Tx (0.1% Triton-X100 [Sigma Aldrich, 9002-93-1] in 1X DPBS) for 5 minutes at RT. After three washes with 1X DPBS, cells were incubated in blocking solution of 2% bovine serum albumin (BSA [Sigma Aldrich, A9647]) prepared in 1X DPBS for 1h at RT. Cells were then incubated with primary antibody overnight (for 16h) at 4°C. The following primary antibodies were used: rabbit anti-EEA1 (Thermo Fisher Scientific, PA1-063A; 1:100), mouse anti-LAMP1 (Developmental Studies Hybridoma Bank, H4A3; 1:100), rabbit anti-LC3 (Medical and Biological Laboratories International Corporation, PM036; 1:400), guinea pig anti-p62 (Progen, GP62-C; 1:400), and rabbit anti-TFEB (Cell Signaling Technology, 4240; 1:100), mouse anti-human Galectin3 (Clone 194804, MAB1154; 1:50), and rabbit anti-human IST1 (Proteintech, 19842-1-AP; 1:100). The next day, cells were washed three times with 0.1% DPBS-Tx, followed by incubation with respective secondary antibodies (all from Thermo fisher Scientific, dilution of 1:500 in blocking solution) for 1h at RT. Cells were then washed three times with 1X DPBS, counterstained for nuclei with DAPI (1:1000 in PBS [Sigma-Aldrich, D9542]) and mounted on glass slides with Aqua-Poly/Mount (Polysciences Inc., 18606-20). Slides were imaged at least a day after mounting of coverslips.

hiPSC-derived cells were fixed with 4% PFA for 20 minutes, followed by three washes with 1X PBS. After three washes with 1X Tris-buffered saline (TBS) and 0.1 % Triton-X100 (Sigma-Aldrich, 9036-19-5), cells were blocked with a solution composed of 0.3% Triton-X100 and 3% normal donkey serum (Millipore, 41105901) diluted in TBS for 2h at RT. Cells were then incubated with primary antibodies in afore-mentioned dilutions overnight at 4°C. The next day, cells were washed three times with 1X TBS and 0.1 % Triton-X100, and incubated with blocking solution for 1h at RT. Secondary antibodies diluted in blocking solution were then added for 2h at RT, followed by three washes with 1X TBS. Finally, cells were incubated with DAPI (Abcam, 228549), diluted to 1:5000 in TBS), and coverslips were mounted on glass slides with PVA-DABCO (Sigma Aldrich) mounting media.

### Lysosome degradative ability and motility assays

To measure the degradative ability of lysosomes, cells were grown on 35 mm glass-bottom microdishes for 24h, and first incubated with AlexaFluor647-conjugated Dextran of 10 kDa molecular weight (Invitrogen, D22914) for 3 hours (pulse). After 3h, cells were washed and replaced with media containing 500 nM *α*-Syn488, or only media for control conditions, and incubated for 16h. DQ-BSA Red (Invitrogen, D12051) was diluted in fresh, complete medium to a final concentration of 10 *μ*g/mL, and added to cells for the last two hours of aggregate incubation at 37°C. Cells were thoroughly washed with complete medium before imaging.

To assess the motility of lysosomes, cells were incubated for 16h with *α*-Syn aggregates conjugated with AlexaFluor 568 fluorophore. After a thorough wash with 1:3 diluted trypsin, cells were incubated with 500 nM Lysotracker Green DND-189 (Thermo Fisher Scientific, L7535) diluted in fresh, complete medium for 30 minutes at 37°C. After a wash with complete medium, fresh phenol red-free RPMI 1640 medium (for SH-SY5Y cells) and FluoroBrite™ DMEM (for HMC3 cells) was added and the dishes were incubated in a 37°C, 5% CO_2_, 50% relative humidity environmental chamber equipped with the microscope. Time-lapse imaging was performed using Nikon Eclipse Ti2 microscope with a spinning disk confocal set-up (Yokogawa) with the following lasers (wavelength in nm): 405, 488, 561 and 640 nm. Cells were imaged using a 100X oil immersion objective (1.45 numerical aperture) for 5 minutes. 5 optical sections of 1 *μ*m interval (± 2 sections from the middle plane) were taken for each frame in two wavelengths – 488 nm and 561 nm. Each acquisition was made at a time interval of 1.537 seconds.

### Autophagy and lysophagy flux analysis

To assess autophagy flux upon exposure of cells to *α*-Syn aggregates, cells were immunostained for p62 (as previously described). For bafilomycin treatment of *α*-Syn containing cells, 400 nM of the drug was added to cells in the fifteenth hour (last 1h of aggregate incubation), followed by fixation. For lysophagy analysis, similar approach was undertaken for aggregate and bafilomycin A1 incubation. Lysophagy was induced by the lysosome-damaging drug L-leucyl-L-leucine methyl ester (LLOMe; Sigma-Aldrich, L7393) at a concentration of 1 mM for 1h, co-incubated with bafilomycin A1. Cells were then washed with 1X DPBS, and processed for immunocytochemistry.

### Immunoblotting

Both SH-SY5Y and HMC3 cells were grown in 6-well dishes for 24h before incubation with untagged *α*-Syn aggregates (500 nM) for 16h. 400 nM Bafilomycin A1 for the last 4 hours before harvesting cells. Following incubation, cells were trypsinized and the pellet fraction was collected. Cell pellets were homogenized in 100 μl of radioimmunoprecipitation assay (RIPA) buffer consisting of 50 mM Tris-HCI (pH 7.4), 1% TritonX-100, 0.5% sodium-deoxycholate, 0.1% sodium dodecyl sulphate, 150 mM sodium chloride and 2 mM ethylenediaminetetraacetic acid, supplemented with 1X protease inhibitor (cOmplete Mini, EDTA-free; Sigma, 11836170001). Protein concentrations were measured using Bradford’s method, following manufacturer’s instructions (Thermo Fisher Scientific). Proteins (20 µg) were denatured in SDS (8%) and β-mercaptoethanol (5%) at 95 °C for 10 minutes. After separation by SDS-PAGE on a 14% gel, proteins were electro-transferred to nitrocellulose membranes. The membranes were blocked with 5% BSA and incubated overnight with primary antibodies (anti-LC3B, D11 XP #3868, Cell Signaling Technology, 1:1000; and anti-GAPDH antibody Sigma, G9545, 1:5000) in blocking buffer, followed by incubation with horseradish peroxidase-conjugated secondary antibody (Millipore, 1:5000) in TBS containing 0.1% Tween-20 at room temperature for 1 h. Finally, proteins were visualized using an ECL kit (Thermo Fisher Scientific) and chemiluminescence images were acquired using GE Amersham Imager AI680 analyzer. ***mRNA isolation and qPCR*** RNA was isolated using TRIZOL reagent (Invitrogen) following manufacturer’s protocol. 1 μg of RNA was used to synthesize cDNA using high capacity cDNA reverse transcription kit (4368814, Applied Biosystems). qPCR was carried out using diluted cDNA (20 ng per reaction) using iTaq universal SYBR Green supermix (1725124, Bio-Rad). Relative mRNA expression was calculated using the ΔΔCt method and each gene was normalized with Ct value of β-actin. Three technical replicate reaction mixes were performed for each gene and biological replicate. Primers against the following human genes were used:

*LAMP1* (forward: CGTGTCACGAAGGCGTTTTCAG; reverse: CTGTTCTCGTCCAGCAGACACT)

*CTSD* (forward: GCAAACTGCTGGACATCGCTTG; reverse: GCCATAGTGGATGTCAAACGAGG)

*CTSB* (forward: GCTTCGATGCACGGGAACAATG; reverse: CATTGGTGTGGATGCAGATCCG)

*ATP6V1H* (forward: CGGGTCAATGAGTACCGCTTTG; reverse: GATACTGGAGCTGAAAGCCACAC)

*SQSTM1* (forward: TGTGTAGCGTCTGCGAGGGAAA; reverse: AGTGTCCGTGTTTCACCTTCCG)

### *α*-Syn transfer assay

To assess for aggregate transfer, neuronal cells or microglia were grown for 24h in a 6-well dish, and then exposed to 500 nM of *α*-Syn aggregates, prepared as previously described. After 16h of incubation, cells (now the donor population) were trypsinized and seeded on 12 mm coverslips. After 22h post-seeding, donor cells were treated (or not) with 200 nM wortmannin (Sigma Aldrich, W1628) for 2h. Cells were then washed with complete media once, and then fresh media was added. At this point, microglia or neuronal cells (acceptor population) were added to the culture (1:1 ratio). For the experiment wherein both donor and acceptor cell populations were treated with wortmannin, the donor population was seeded first on coverslips, and grown for 22h. Media was replaced and the acceptor population was added, along with wortmannin. Media was replaced after 2h of wortmannin treatment. Co-cultures were done for 12h, after which cells were fixed to preserve TNTs (described below) and stained with wheat germ agglutinin 647 (WGA, Thermo Fisher Scientific, W32466; 3.33 *μ*g/mL) for 15 minutes at RT, followed by nuclei counterstaining with DAPI (1:1000 in 1X DPBS).

To assess the extent of secretion-mediated aggregate transfer, conditioned media from aggregate containing cells treated (or not) with wortmannin was added to acceptor microglia for 12h. Data represented is normalized to secretion control (data from secretion control subtracted from the co-culture data).

### TNT counting

To efficiently visualize TNTs in cultures, cells were fixed at sub-confluency (∼70%). In order to appropriately preserve TNTs, two different fixative solutions were used, as described previously (Chakraborty *et al*, 2023b; Sáenz-de-Santa-María *et al*, 2023): fixative 1 (0.05% glutaraldehyde [GA {Sigma Aldrich, G5882}], 2% PFA, 0.2 M HEPES buffer [Gibco, 15630-080] in 1X DPBS), followed by fixative 2 (4% PFA, 0.2 M HEPES buffer in 1X DPBS) for 15 minutes each at RT. Cells were then labelled with Phalloidin 647 (Thermo Fisher Scientific, A12380; 1:250 in 1X DPBS) and DAPI (1:1000 in 1X DPBS) for 15 minutes each at RT. Cells were then washed once with 1X DPBS and mounted on glass slides.

### Fixed-cell microscopy

Images were acquired using Zeiss LSM900 inverted confocal microscope equipped with four lasers (wavelength in nm): 405, 488, 561 and 640 nm. For TNT counting, samples were imaged using 40X oil immersion objective (1.3 numerical aperture) with a field-of-view effective zoom of 0.8x, whereas all other immunofluorescence samples were acquired using a 63X oil immersion objective (1.4 numerical aperture) with a 1x zoom. Image acquisition was performed using ZEN blue software. The entire cell volume was imaged for all samples, with optical sections of 0.45 *μ*m. Depending on the cell types, the entire volume of cells ranged between 7 and 13 *μ*m in thickness.

Super-resolution images were acquired using Zeiss LSM 780 Elyra SIM set-up (Carl Zeiss, Germany) using Plan-Apochromat 63×/1.4 oil objective with a 1.518 refractive index oil (Carl Zeiss). 16-bit images were acquired in “frame-fast” mode between wavelengths, with 32.0 *μ*m grid size. Optical thickness was set at 0.133 *μ*m. Raw images were processed using the SIM processing tool of Zen black software.

### Quantification and Statistical Analyses

For TNT counting, images were processed for analysis using the “manual TNT annotation” plug-in of ICY software (https://icy.bioimageanalysis.org/plugin/manual-tnt-annotation/). Number of p62 puncta or *α*-Syn aggregates per cell were manually counted in 3D images, through the z-stacks. Colocalization analysis was performed using the JACoP plug-in FIJI (Bolte & Cordelières, 2006). 3D images were used for colocalizing two channels with object-based detection, and thresholding of objects was performed manually. For analyzing degradative abilities of lysosomes, selections were created from dextran channel, and intensity of DQ-BSA was measured in all lysosomes. To measure DQ-BSA intensity in lysosomes positive for *α*-Syn, “image calculator” function of FIJI was used, to identify pixels positive for both dextran and *α*-Syn (AND operation). The resultant image was thresholded and made to a selection, followed by masking on the DQ-BSA channel before measuring the intensity. Triple colocalization events were counted manually for each cell in 3D images. Time-lapse videos were analyzed for ∼2 minutes (80 frames) to track lysosomal motility. Manual tracking was performed using the FIJI plug-in MTrackJ (Meijering *et al*, 2012) for single lysosomes. Only those lysosomes were tracked that were not in the perinuclear area, and that remained within the region of interest for the entire duration of tracking without merging/interacting with other lysosomes. Movies were then created at a frame speed of 5 fps.

No prior power analysis was done to measure the sample size. Graphs were plotted using GraphPad prism 10.0, and appropriate statistical tests were performed on raw data. For all datasets, an initial normality distribution test was performed, and non-parametric tests were performed for any datasets that did not satisfy normal distribution. Statistical tests performed are mentioned in the figure legends. All the experiments were performed for three independent biological replicates.

## Supporting information

Supplementary Figures with legends, and Movie legends

## Acknowledgments

We thank Dr. Stephanie Maya and Dr. Valeria Valente for technical assistance with experiments. We thank Dr. Christel Brou, and all the members of the Membrane Traffic and Pathogenesis Unit, Institut Pasteur, for insightful discussions, Dr. Francesca Palese and Sevan Belian for critical reading of the manuscript. We thank Reine Bouyssie, a member of the administrative staff of the Membrane Traffic and Pathogenesis Unit for her continued support.

## Funding

Pasteur-Paris University International Doctoral Program (RC)

The Journal of Cell Science Travelling Fellowship JCSTF24101615 (RC)

Biology Summer Studentship Programme - Institut Pasteur and Trinity College, University of Cambridge (PS)

France Parkinson - Soutien de l’Association France Parkinson 2021 (CZ)

Don Explore AD - Programme Explore de l’Institut Pasteur (CZ)

Agence Nationale de la Recherche ANR-20-CE13-0032-01 (CZ)

Fondation pour la Recherche Médicale FRM - EQU202103012692 (CZ)

PID2022-139546OB-I00 supported by MCIN/AEI/10.13039/501100011033 and FEDER, and PDC2021-121051-I00 supported by MCIN/AEI/10.13039/501100011033 and by the European Union Next Generation EU/ PRTR) (AC)

AGAUR (2021-SGR-974) (AC)

VT was the recipient of a pre-doctoral La Caixa INPhINIT Incoming Fellowship (code: LCF/BQ/DI21/11860038

JMM was the recipient of a pre-doctoral fellowship FPI (PRE2022-104573) from the Spanish Ministry of Economy and Competitiveness (MINECO).

Research was conducted within the context of Pasteur International Joint Research Unit Neurodegenerative Diseases.

## Author contributions

Conceptualization: CZ, RC; Methodology: RC, PS, VT, JMM, TN, MH; Investigation: RC, PS, VT, JMM; Visualization: RC; Supervision: CZ, AC; Funding acquisition: CZ, AC; Writing—original draft: RC; Writing—review & editing: RC, PS, VT, JMM, AC, CZ.

## Data and materials availability

All data are available in the main text, or in the supplementary materials.

