## Supplementary Figures with legends, and Movie legends for "Neuronal Autophagy Failure Drives α-Synuclein Transfer to Microglia to Outsource Aggregate Clearance"

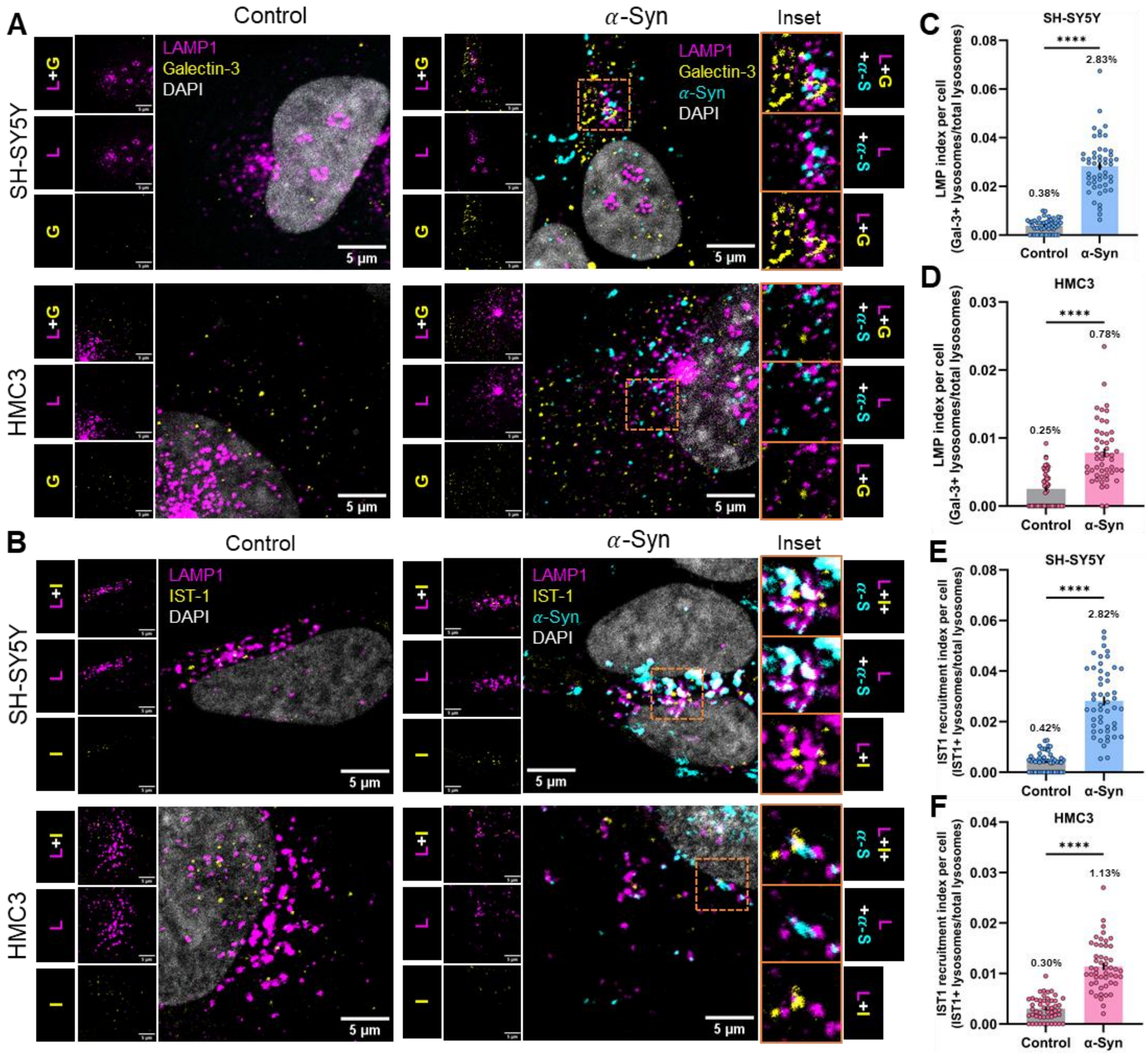


**Figure S1.** Lysosomal damage in neuronal and microglial cells. (**A**) Representative images of SH-SY5Y cells (upper panels) and HMC3 cells (bottom panels) immunostained for Galectin-3 (yellow) and LAMP1 (magenta) in control and $\boldsymbol{\alpha}$-Syn (cyan) treated conditions. Insets show regions of triple colocalization of $\boldsymbol{\alpha}$-Syn-associated lysosome with its membrane permeabilized. (**B**) Representative images of SH-SY5Y cells (upper panels) and HMC3 cells (bottom panels) immunostained for IST-1 (yellow) and LAMP1 (magenta) in control and $\boldsymbol{\alpha}$-Syn (cyan) treated conditions. Insets show regions of triple colocalization of $\boldsymbol{\alpha}$-Syn-associated lysosome with recruited ESCRT-III related protein IST1 for potential membrane repair. (**C and D**) Quantification of the percentage of lysosomes per cell with membrane permeabilized in neuronal cells (**C**) and microglia (**D**). Mean percentage is mentioned within the graph. Error bars represent SEM. N=3 independent experiments, n=50 cells per group. Statistical significance was analyzed using Mann-Whitney U test. ****p<0.0001. (**E and F**) Quantification of the percentage of lysosomes per cell with IST1 recruitment in neuronal cells (**E**) and microglia (**F**). Mean percentage is mentioned within the graph. Error bars represent SEM. N=3 independent experiments, n=50 cells per group. Statistical significance was analyzed using Mann-Whitney U test. ****$p<0.0001$.

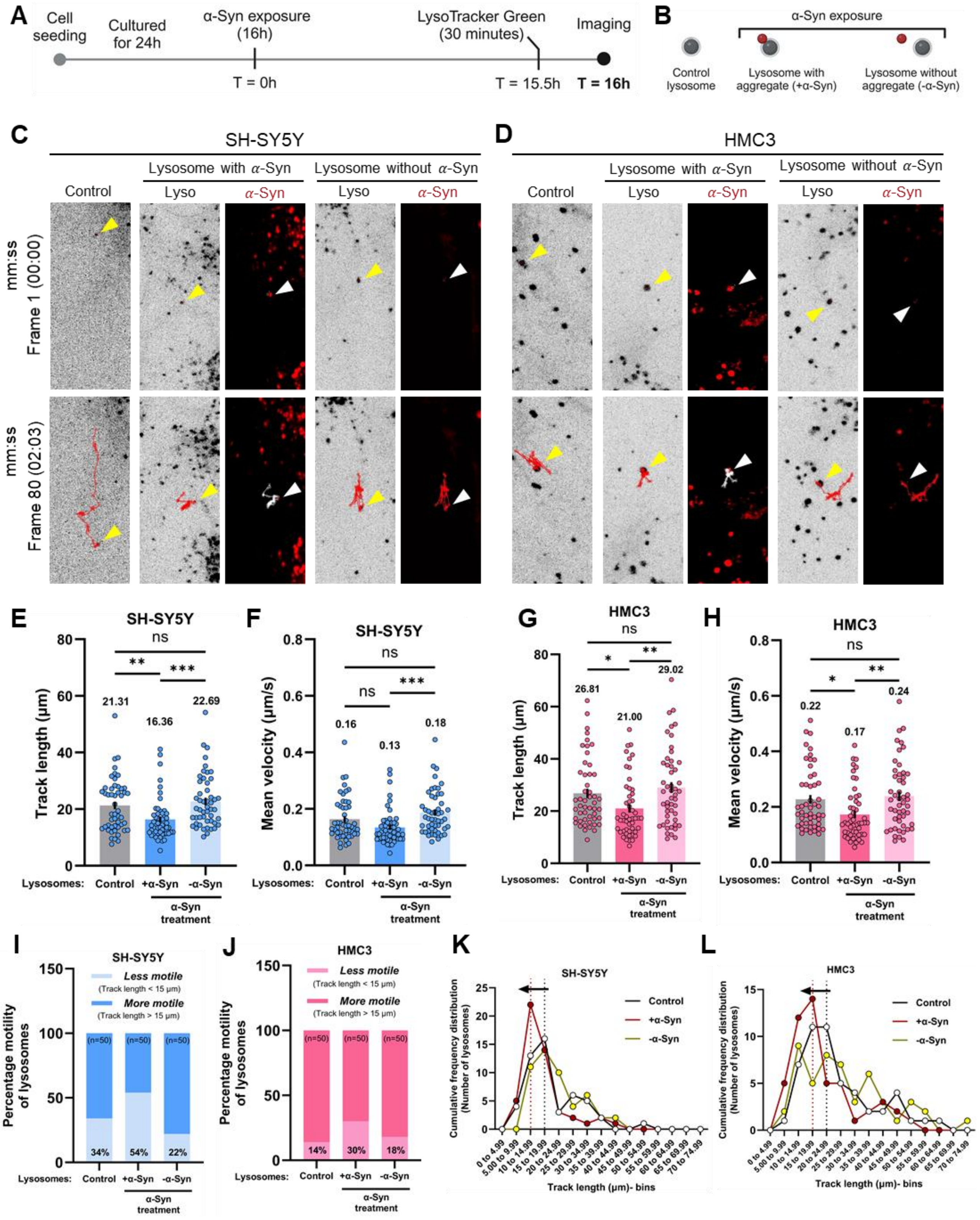


**Figure S2.** (**A and B**) Schematic representation of the experimental (**A**), and analysis strategy (**B**) to track lysosome motility. (**C and D**) Representative images of the first frame (upper panels) and last (80th) frame (lower panels) of tracked lysosomes for SH-SY5Y cells (**C**) and HMC3 cells (**D**). Arrowheads point towards the tracked lysosome. (**E**) Mean track lengths of lysosomes in neuronal cells. Statistical significance was analyzed using Kruskal-Wallis test with Dunn's multiple comparison. ns: non-significant ($p>0.9999$),

**p=0.0067, ***p<0.001. (**F**) Mean velocities of lysosomes in neuronal cells. Statistical significance was analyzed using Kruskal-Wallis test with Dunn's multiple comparison. ns: non-significant (p=0.0664 between control and +$\alpha$-Syn groups, p=0.2038 for control and -$\alpha$-Syn groups), ***p=0.0001. (**G**) Mean track lengths of lysosomes in microglia. Statistical significance was analyzed using Kruskal-Wallis test with Dunn's multiple comparison. ns: non-significant (p>0.9999), *p=0.0319, **p=0.0052. (**H**) Mean velocities of lysosomes in microglia. Statistical significance was analyzed using Kruskal-Wallis test with Dunn's multiple comparison. ns: non-significant (p>0.9999), *p=0.0182, ***p=0.0052. (**I and J**) Proportion of lysosomes that travel less than 15 $\mu$m (less motile), or more than 15 $\mu$m (more motile) in different conditions for neuronal cells (**I**) and microglia (**J**). For quantifications in (**E-J**): N=3 independent experiments, n=50 individual lysosomes tracked per group. Mean values are mentioned within the graphs. Error bars represent SEM. (**K and L**) Cumulative frequency distribution of the 50 tracked lysosomes of SH-SY5Y (**K**) and HMC3 (**L**) cells that travel different distances, binned in clusters of 4.99 μm. Black vertical line indicates the peak point for control lysosomes, and red vertical line indicates the same for lysosomes associated with α-Syn. Black arrows within the graphs indicate a shift in the peak (less distance travelled by aggregate-associated lysosomes).

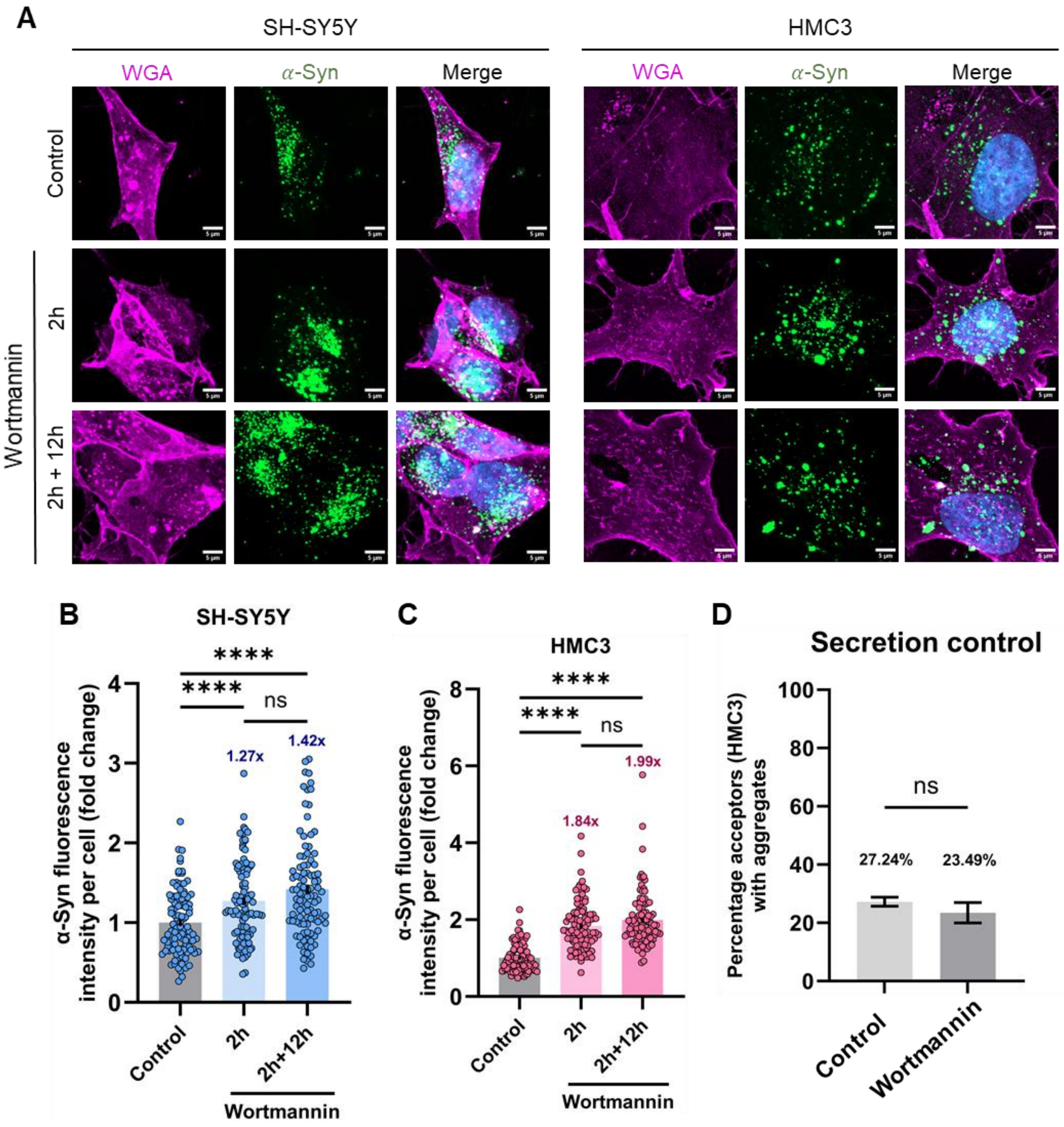


**Figure S3.** Load of aggregates in neuronal cells and microglia upon wortmannin treatment. (**A**) Representative images of SH-SY5Y cells (left panels) and HMC3 cells (right panels) after 16h of exposure to $\alpha$-Syn aggregates (control, upper panels), followed by 2 h treatment with wortmannin (2 h, middle panels), and grown for additional 12 h after washing wortmannin off (2 h + 12 h, lower panels). (**B and C**) Quantification of intracellular aggregate load in different conditions for SH-SY5Y (**B**) and HMC3 (**C**) cells. Data represented as fold change relative to control. Fold difference is mentioned within the graph. N=3 independent experiments, n=100 cells per condition. Error bars represent SEM. Statistical significance was analyzed using Brown-Forsythe and Welch One-Way ANOVA with Games-Howell's multiple comparison. ns: non-significant (p=0.1548 in neuronal analysis, and p=0.2513 in microglial analysis), ****p<0.0001. (**D**) Secretion control of aggregate transfer from neuronal cells (wortmannin treated or not) to microglia.

Data of percentage of acceptor cells positive for $\alpha$-Syn aggregates represented as mean ± SEM. N=3 independent experiments, number of acceptor cells analyzed (n)= 191 cells for control and 259 cells for wortmannin-treated condition. Statistical significance was measured using unpaired Student's t-test with Welch's correction. ns: non-significant.

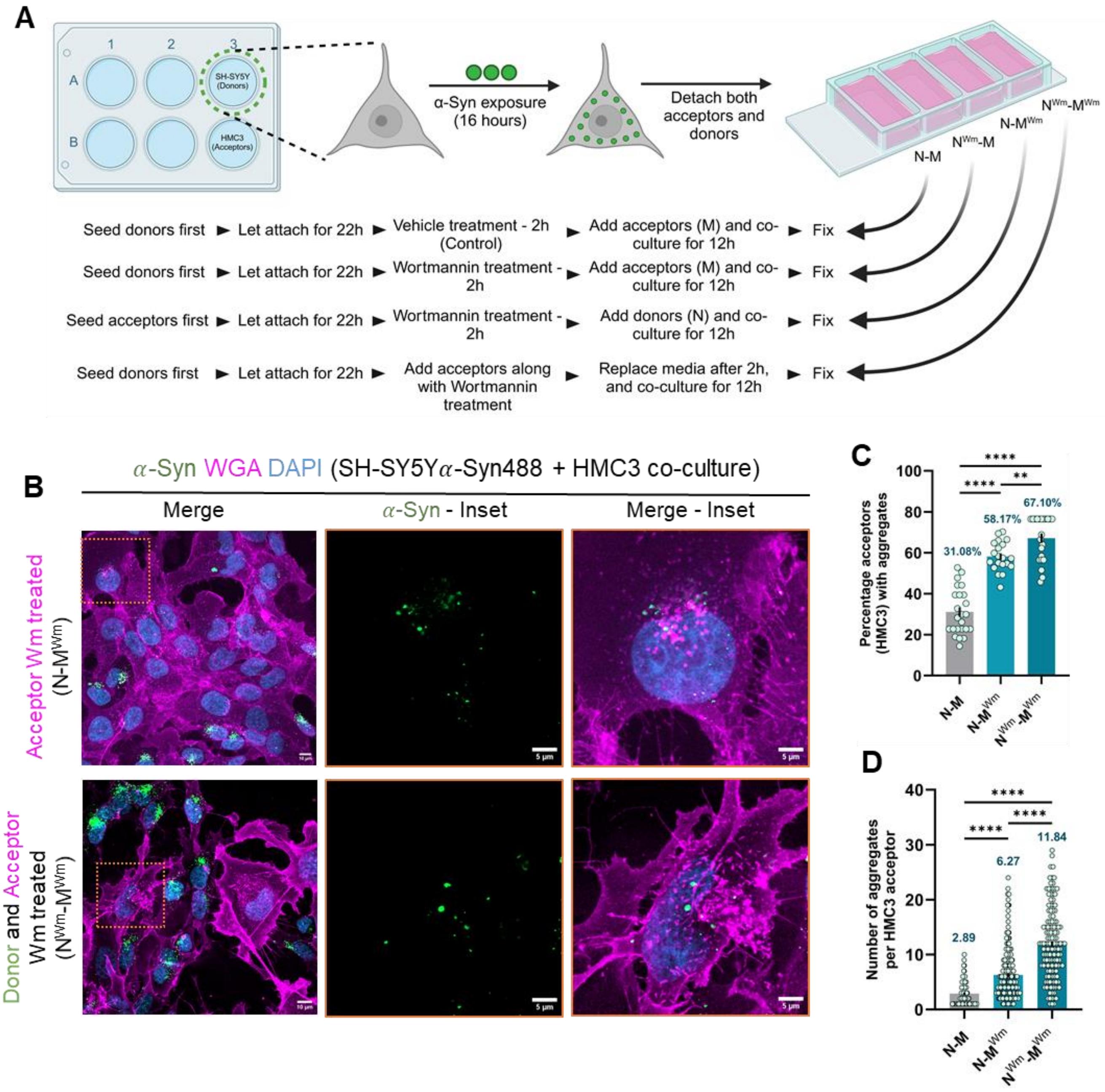


**Figure S4.** Contribution of autophagy in transfer of aggregates, and its relation to TNTs. (**A**) Schematic representation of co-culture strategy. (**B**) Representative images of 12h co-cultures between $\alpha$-Syn loaded SH-SY5Y neuronal cells and HMC3 microglia (wortmannin treated in different combinations. Wortmannin treatments of respective cell populations are represented by the superscript "Wm". Dotted box indicates the region of acceptor microglia zoomed in. (**C**) Percentage of HMC3 acceptor cells that received aggregates in control and wortmannin-treated cocultures. Average percentage of acceptor cells positive for aggregates is mentioned within the graph. N=3 independent experiments, n=240 acceptor cells for N-M (same as in Figure 5A-B), 322 acceptor cells for N-$M^{Wm}$, and 161 acceptor cells for $N^{Wm}$-$M^{Wm}$. Error bars represent SEM. Statistical significance was analyzed using Brown-Forsythe and Welch One-Way ANOVA with Dunnett's T3 multiple comparison. ****p<0.0001 (**D**) Quantification of the number of aggregates received per cell by acceptor HMC3 in (**B**). Mean number of aggregates is mentioned within the graph. N=3 independent experiments. N=137 acceptor cells for N-M (same as in Figure 5A, C), 264 acceptor cells for N-$M^{Wm}$, and 144 acceptor cells for $N^{Wm}$-$M^{Wm}$. Error bars represent SEM. Statistical significance was analyzed using Kruskal-Wallis test with Dunn's multiple comparison. ****p<0.0001.

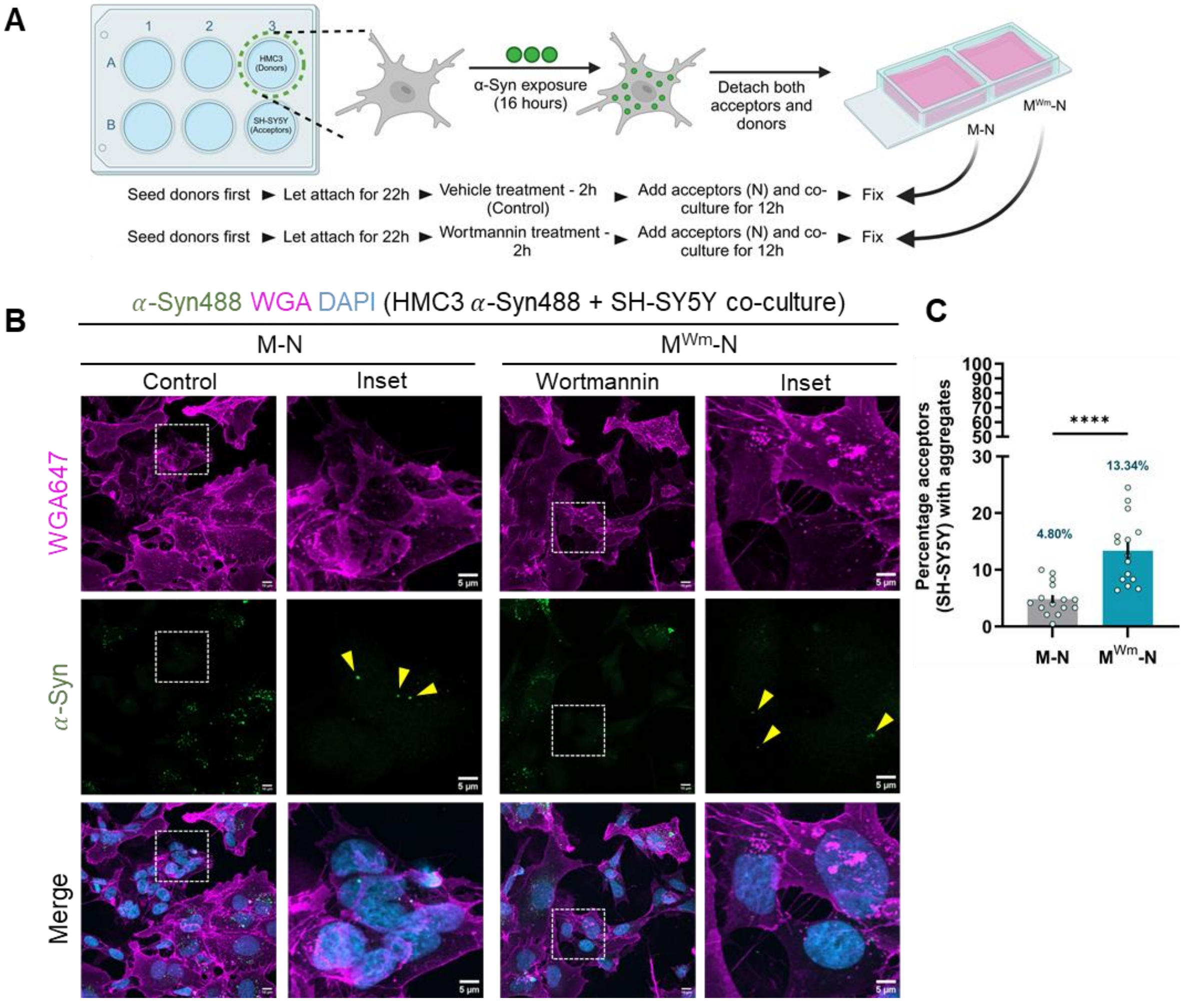


**Figure S5.** Contribution of autophagy impairment in aggregate transfer. (**A**) Schematic of co-culture experimental design to assess aggregate transfer from microglia to neuronal cells upon autophagy inhibition by Wm. (**B**) Representative images of co-culture between ***α***-Syn loaded HMC3 cells (control – left panels and wortmannin treated – right panels) and SH-SY5Y neuronal cells. Dotted box indicates the region zoomed in. Yellow arrowheads point towards the aggregates. (**C**) Percentage of SH-SY5Y acceptor cells that received aggregates in control (M-N) and wortmannin-treated HMC3 ($M^{Wm}$-N) co-cultures. Average percentage of acceptor cells positive for aggregates mentioned in red within the graph. N=3 independent experiments, n=322 acceptor cells for control and 327 acceptor cells for wortmannin treated groups. Error bars represent SEM. Statistical significance was analyzed using unpaired Student's t-test with Welch's correction. ****p<0.0001.

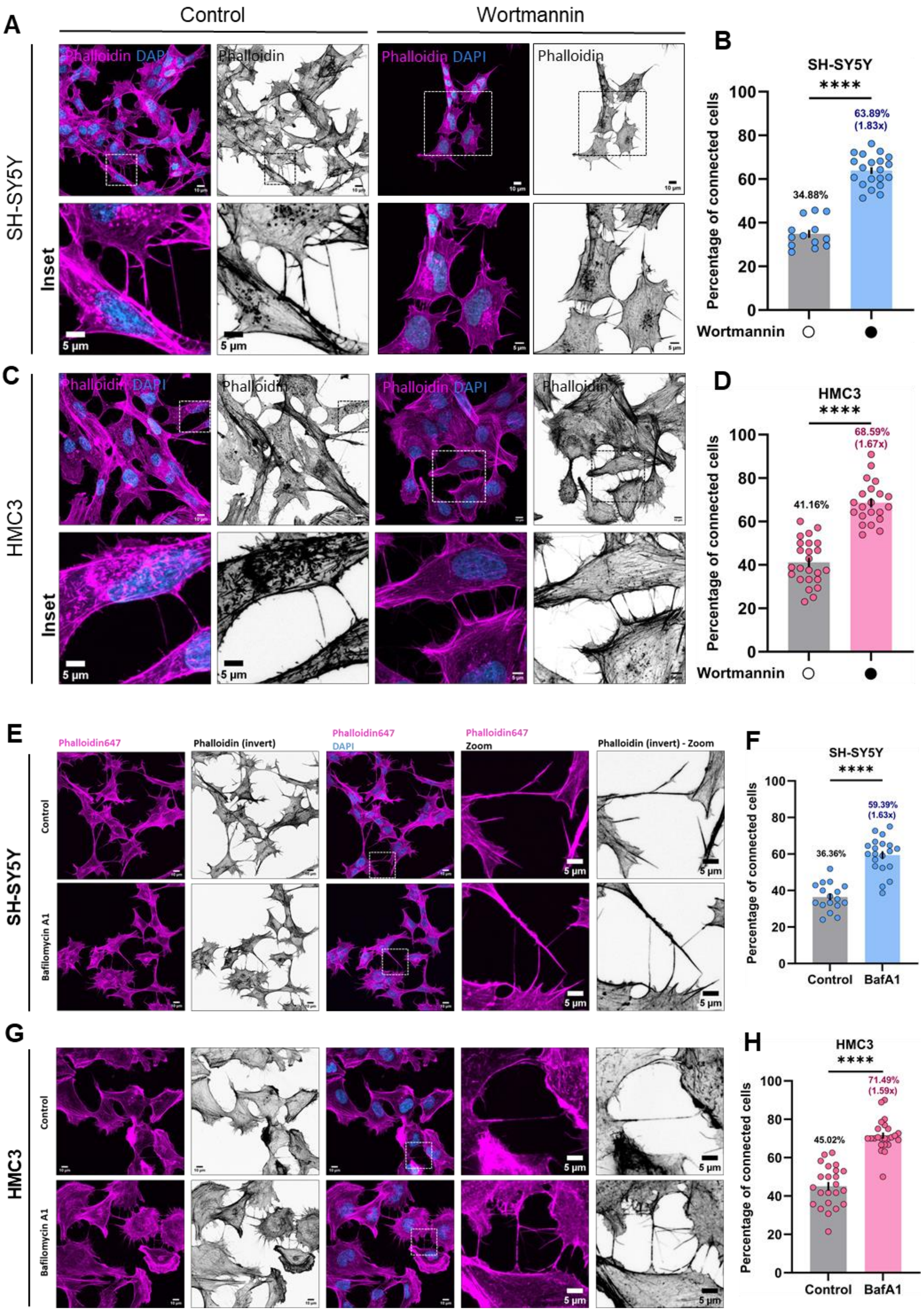
A
Control
Wortmannin
Phalloidin DAPI
Phalloidin
SH-SY5Y
Inset
5 µm
B
SH-SY5Y
****
63.89%
(1.83x)
34.88%
Percentage of connected cells
Wortmannin
C
HMC3
D
HMC3
68.59%
(1.67x)
41.16%
E
Phalloidin647
Phalloidin (invert)
DAPI
Zoom
Phalloidin (invert) - Zoom
Control
Bafilomycin A1
F
59.39%
(1.63x)
36.36%
BafA1
G
H
71.49%
(1.59x)
45.02%

**Figure S6**: Autophagy inhibition and intercellular connections. (**A and C**) Representative images of control (left panels) and wortmannin-treated (right panels) SH-SY5Y (**A**) and HMC3 (**C**) cells, stained for phalloidin (magenta, and gray-invert LUT). Dotted box indicates the region zoomed in. (**B**) Quantification of the percentage of connected SH-SY5Y cells. Mean percentage and fold difference is mentioned within the graph. N=3 independent experiments, n=551 cells for control, and 641 cells for wortmannin treated groups. (**D**) Quantification of the percentage of connected HMC3 cells. Mean percentage and fold difference is mentioned within the graph. N=3 independent experiments, n=368 cells for control, and 275 cells for wortmannin treated groups. For both (**B**) and (**D**), error bars represent SEM. Statistical significance was analyzed using unpaired Student's t-test with Welch's correction. ****p<0.0001. (**E**) Representative images of SH-SY5Y cells treated or not with bafilomycin A1 and stained with phalloidin to assess TNTs. (**F**) Quantification of the percentage of TNT-connected SH-SY5Y cells. Fold difference mentioned in red within the graph. N=3 independent experiments, n=398 cells for control and 500 cells for bafilomycin A1 treated condition. Statistical significance was analyzed using unpaired Student's t-test with Welch's correction. ****p<0.0001. (**G**) Representative images of HMC3 cells treated or not with bafilomycin A1 and stained with phalloidin to assess TNTs. (**H**) Quantification of the percentage of TNT-connected HMC3 cells. Fold difference is mentioned within the graph. N=3 independent experiments, n=205 cells for control and 215 cells for bafilomycin A1 treated condition. Statistical significance was analyzed using unpaired Student's t-test with Welch's correction. ****p<0.0001.

### Supplementary movie legends

**Movie S1.** Tracking lysosomal movement in control condition of SH-SY5Y cells.

**Movie S2.** Tracking lysosomal movement upon $\alpha$-Syn treatment of SH-SY5Y cells. Representative time-lapse movie of lysosome containing aggregates.

**Movie S3.** Tracking lysosomal movement upon $\alpha$-Syn treatment of SH-SY5Y cells. Representative time-lapse movie of lysosome without aggregates.

**Movie S4.** Tracking lysosomal movement in control condition of HMC3 cells.

**Movie S5.** Tracking lysosomal movement upon $\alpha$-Syn treatment of HMC3 cells. Representative time-lapse movie of lysosome containing aggregates.

**Movie S6.** Tracking lysosomal movement upon $\alpha$-Syn treatment of HMC3 cells. Representative time-lapse movie of lysosome without aggregates.
